# Tissue-specific regulation of PNPLA3 promotes lipid remodeling in response to dietary and environmental challenges

**DOI:** 10.1101/2025.10.27.684800

**Authors:** Panyun Wu, Yang Wang, Jonathan C. Cohen, Helen H. Hobbs

## Abstract

**Background & Aims:** PNPLA3(I148M) is the strongest genetic risk factor for steatotic liver disease (SLD), but its functional role and tissue-specific regulation remain unclear. In mice, PNPLA3 is abundant in liver, yet undetectable in adipose depots. Here, we characterize the molecular mechanisms underlying these tissue-specific differences in PNPLA3 expression in mice to clarify its functional role and link to SLD risk.

**Methods:** *Pnpla3* mRNA and PNPLA3 protein levels were quantified in liver and adipose depots of fasted and refed mice at 30°C and 6°C. Signaling pathways regulating PNPLA3 expression in adipocytes were examined using adrenergic agonists and pathway-specific modulators. Translation and proteasomal inhibitors were used during adrenergic stimulation to investigate the discordance between *Pnpla3* mRNA and protein levels. Relationship between PNPLA3 levels and triglyceride (TG) fatty acid composition was also assessed.

**Results:** At thermoneutrality, feeding strongly increased PNPLA3 levels in liver but it remained undetectable in adipose tissue of mice. Conversely, cold exposure or β3-adrenergic stimulation had no effect on hepatic PNPLA3, but increased PNPLA3 >19-fold in brown adipose tissue (BAT), despite causing a >75% reduction in *Pnpla3* mRNA, indicating robust post-translational regulation. In BAT, adrenergic signaling via cAMP/PKA and PI3K/AKT elevated PNPLA3 by reducing proteasomal degradation. PNPLA3 expression correlated with depletion of TG-long-chain polyunsaturated fatty acids (TG-LCPUFAs) in both liver and BAT, consistent with a role in lipid remodeling.

**Conclusions:** These findings reveal striking tissue- and context-specific regulation of PNPLA3, but a conserved association between its expression and TG-LCPUFAs levels, suggesting that PNPLA3 modulates lipid remodeling in response to metabolic stress and that disrupting this function may contribute to SLD susceptibility.

**Impact and implications:** Despite being the strongest genetic risk factor for SLD, PNPLA3’s physiological role remains unclear. Using mouse models, this study reveals that PNPLA3 is regulated in a tissue-specific manner in response to feeding and cold exposure, thereby promoting remodeling of cellular lipids to adapt to dietary and environmental challenges. The localization of PNPLA3 action and its tissue-specific regulation are directly relevant to hepatologists and metabolic researchers aiming to understand its influence on intracellular lipid composition and its effects on disease susceptibility. Moreover, modulation of PNPLA3 turnover—and its impact on LCPUFAs remodeling—emerges as a potential therapeutic strategy for regulating lipid homeostasis in SLD.

**Graphical Abstract:** 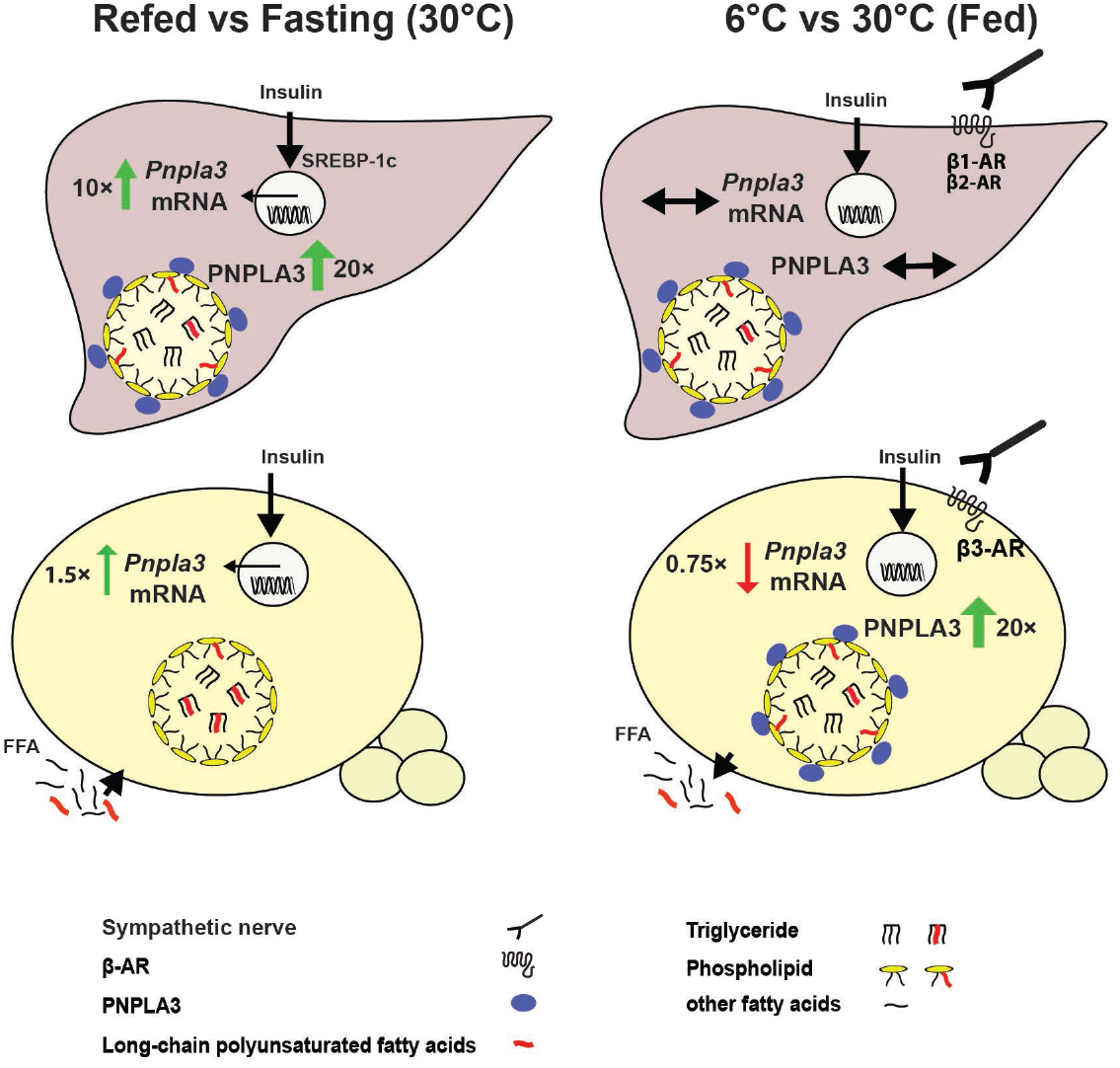

**Highlights:**

- PNPLA3 is regulated in a highly tissue-specific manner in mice.
- In liver, feeding–but not cold exposure–induces PNPLA3 primarily through transcriptional mechanisms.
- In adipose tissue, cold exposure–but not feeding–induces PNPLA3 through post-transcriptional mechanisms.
- In adipose tissue, cold exposure increases PNPLA3 despite a reduction in *Pnpla3* mRNA.
- PNPLA3 remodels lipids in liver and adipose tissue to maintain lipid homeostasis, a process disrupted in SLD.

## Introduction

A missense variant (I148M) in patatin-like phospholipase domain-containing protein 3 (PNPLA3) is the strongest genetic risk factor for steatotic liver disease (SLD)^[1]^. The 148M variant is associated with increased hepatic triglyceride (TG) content and progressive liver disease in metabolic dysfunction-associated and alcohol-associated SLD^[1, 2]^. Despite extensive study, the physiological function of PNPLA3 and its relationship to SLD pathogenesis remain unclear.

PNPLA3 shares structural similarity with the principal tissue TG lipase, adipose triglyceride lipase (ATGL)^[3]^, and exhibits selectivity for long-chain polyunsaturated fatty acids (LCPUFAs) over monounsaturated or saturated TG-fatty acids (FAs)^[4, 5]^. Although PNPLA3 enzymatic activity has been characterized, its physiological role is not established. *Pnpla3* is highly expressed in liver and adipose tissue^[6]^. In liver, PNPLA3 is robustly upregulated by feeding through transcriptional activation by the insulin-responsive transcription factor sterol regulatory element-binding protein-1c (SREBP-1c)^[7]^. Hepatic PNPLA3 is stabilized on lipid droplet (LD)^[7]^ and degraded by the ubiquitin-proteasome system^[8, 9]^.

In human adipose tissue, PNPLA3 is expressed at higher levels than in liver^[10]^. In mice, PNPLA3 expression in adipose tissue has not been well-characterized due to lack of reliable antibodies. Therefore, PNPLA3 protein expression has been monitored indirectly, using *Pnpla3* mRNA levels. In mouse adipose tissue, *Pnpla3* mRNA levels change more modestly with food intake than in liver^[6, 7]^, and decrease with cold exposure or β-adrenergic stimulation^[11]^. Despite *Pnpla3* mRNA levels increasing dramatically with adipocyte differentiation^[6]^, the protein is dispensable for adipose tissue formation^[12, 13]^.

To further define the metabolic role of PNPLA3, we compared *Pnpla3* mRNA and protein levels in fat depots and liver. At baseline, *Pnpla3* mRNA levels were 8-fold higher in adipose tissues than liver, yet PNPLA3 was undetectable by immunoblotting. Interestingly, cold exposure markedly increased PNPLA3 protein levels, especially in brown adipose tissue (BAT), but not in liver. The increase in PNPLA3 in BAT was associated with a dramatic decrease in *Pnpla3* mRNA levels. The striking differences in regulation of PNPLA3 in liver and adipose tissue inform the relative roles of the enzyme in lipid metabolism in these two tissues and may be linked to the adverse consequences of genetic variation in *PNPLA3*.

## Materials and methods

The materials and methods are detailed in the *Supplementary Data*.

## Results

### Dissociation between levels of *Pnpla3* mRNA and protein in adipose tissue

*Pnpla3* mRNA and protein levels were compared in liver, BAT, subcutaneous and visceral white adipose tissue (SQ-WAT and V-WAT, respectively) of wild-type (WT) and PNPLA3 148M knockin (*Pnpla3*^*M/M*^)^[14]^ mice maintained at thermoneutrality (30°C) on a high-sucrose diet (HSD)^[7]^. *Pnpla3* mRNA levels were significantly higher in all adipose depots than in liver, with the highest expression in BAT, followed by SQ-WAT and V-WAT, as reported previously in mice maintained at room temperature^[6]^; levels did not differ between WT and *Pnpla3*^*M/M*^ mice (Fig. 1A, left). Despite the higher levels of *Pnpla3* mRNA, PNPLA3 was not detected in adipose tissue lysates, whereas PNPLA3 was readily detected in hepatic LDs (Fig. 1A, right). Consistent with prior findings, PNPLA3 levels in hepatic LDs were higher in *Pnpla3*^*M/M*^ mice than in WT mice^[8]^.

**Fig. 1.**
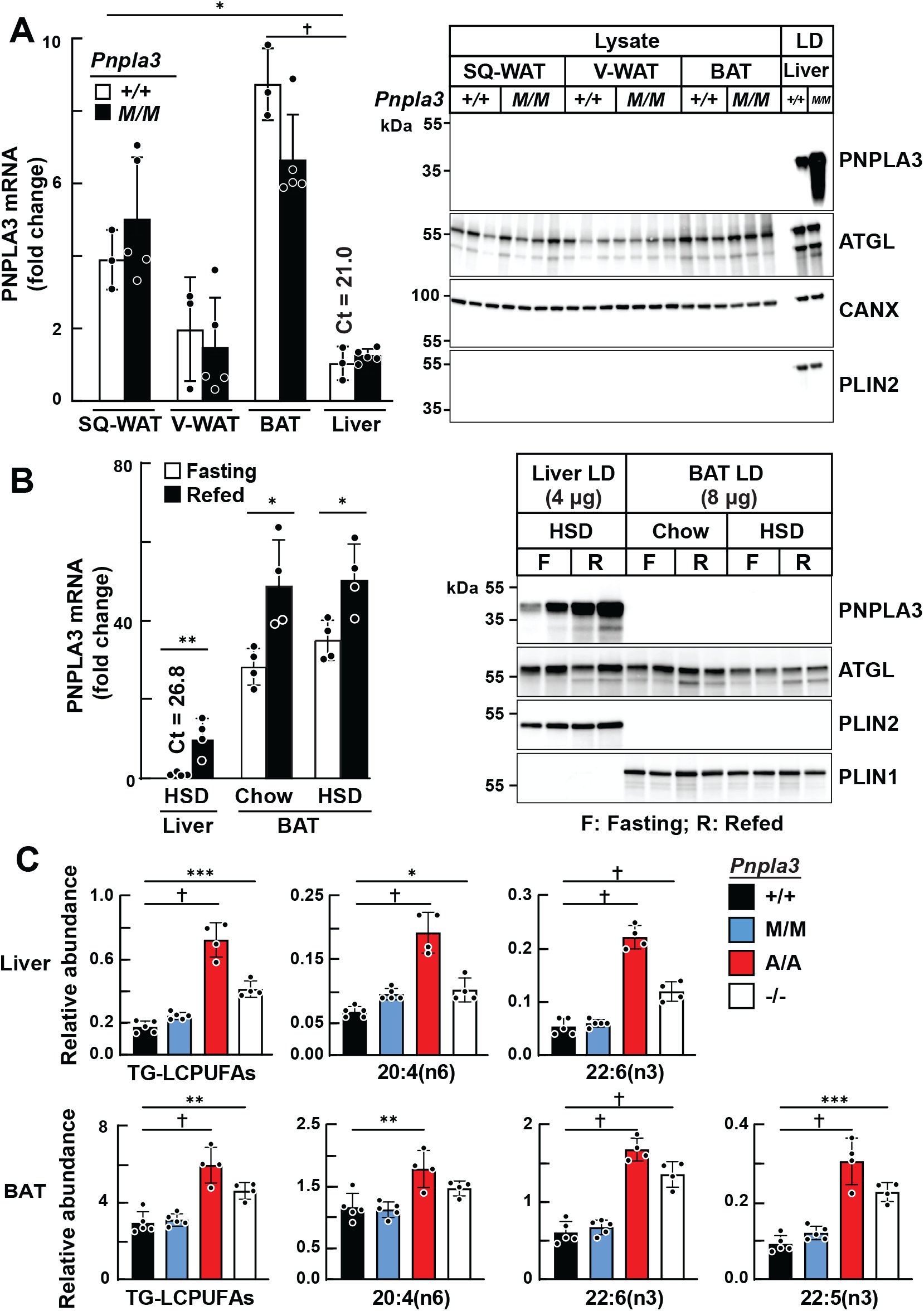
Dissociation between *Pnpla3* mRNA and protein levels in adipose depots. (A) *Pnpla3* mRNA levels in adipose depots and liver, and PNPLA3 immunoblot of adipose lysates (26 µg) and hepatic LDs (6 µg) from WT and *Pnpla3*^*M/M*^ male mice (n=3–5/group) maintained at 30°C. (B) *Pnpla3* mRNA and PNPLA3 protein levels in liver and BAT of WT male mice (n=4/group) under fasting or refeeding conditions on chow or high-sucrose diet. PNPLA3 was assessed in LDs from liver and BAT. (C) TG-LCPUFA profiles in liver and BAT of *Pnpla3*^*-/-*^, *Pnpla3*^*M/M*^, *Pnpla3*^*A/A*^, and WT mice (n=4–5/group) maintained at 30°C. Data represent mean ± SD. P values were determined by Student’s *t*-test (A, B) and by one-way ANOVA followed by Tukey’s multiple comparisons test (C); ^*^P < 0.05; ^**^P < 0.01; ^***^P < 0.001; †P < 0.0001. CANX = calnexin; C_t_ = cycle threshold.

The experiment was repeated using LDs from BAT. In liver, *Pnpla3* mRNA was low after a 12-h fast and increased 11-fold with refeeding^[9]^. In BAT, *Pnpla3* mRNA was 36-fold higher than in liver under fasting conditions and increased only 1.5-fold upon refeeding (Fig. 1B, left). Nevertheless, PNPLA3 remained undetectable in BAT LDs, whereas other LD proteins were readily detected (Fig. 1B, right).

### Fatty acid profiles of BAT resemble those of liver in genetically modified mice

To assess PNPLA3 activity in adipose tissue by an orthogonal approach, we measured levels of TG-LCPUFAs in adipose tissue. If PNPLA3 is expressed in adipose tissue, we would expect the enrichment pattern of TG-LCPUFAs in adipose tissue to resemble that in liver. As observed previously^[4]^, TG-LCPUFAs were depleted in *Pnpla3*^*M/M*^ and WT mice and enriched in *Pnpla3*^*A/A*^ and *Pnpla3*^*-/-*^ mice (Fig. 1C, top). A similar genotype-specific enrichment pattern of TG-LCPUFAs was found in BAT (Fig. 1C, bottom), V-WAT and SQ-WAT (Fig. S1A), whereas other TG-FAs (C16:0, C18:1, C18:2, and C18:3) did not differ significantly among genotypes (Fig. S1B). Previously, we showed in liver that levels of phospholipid (PL)-LCPUFAs are reciprocally related to levels of TG-LCPUFAs^[4]^. Attempts to measure PL-LCPUFAs in BAT LDs were confounded by mitochondrial contamination during LD isolation, a problem encountered previously^[15]^.

The concordant genotype-specific changes in TG-LCPUFAs in liver and BAT suggested that PNPLA3 is active in BAT despite our inability to detect it (Fig. 1). To determine if TG remodeling in BAT was driven indirectly by hepatic PNPLA3, PNPLA3 was overexpressed in livers of *Pnpla3*^*-/-*^ mice using a recombinant adenovirus (Ad-PNPLA3). PNPLA3 was readily detected in liver but remained undetectable in BAT (Fig. S2A). Baseline TG-LCPUFA levels in tissues of *Pnpla3*^*-/-*^ mice were ∼2-fold higher than those in WT mice (Fig. S2B). Hepatic overexpression of PNPLA3 normalized liver TG-LCPUFA levels but did not alter TG-LCPUFA levels in BAT or V-WAT (Fig. S2B). No significant differences in other FA species were observed in TGs from these tissues (Fig. S2C). We concluded that PNPLA3 is expressed in BAT, albeit at levels below the detection limit of our assay. Alternatively, PNPLA3 may be expressed intermittently in BAT, and absent at the time of tissue sampling.

### Cold exposure increases PNPLA3 while reducing *Pnpla3* mRNA in adipose tissue

Given that the highest levels of *Pnpla3* mRNA expression were in BAT, which plays a key role in thermogenesis^[16]^, we examined the effect of cold exposure on PNPLA3 expression in BAT. Mice were housed at 22°C under a 12-h fasting (light)/12-h refeeding (dark) cycle (Fig. 2A). Mice were then maintained at 30°C, 22°C, or 6°C, with half the mice fasted for 12 h and the other half fasted for 6 h and then refed for 6 h. PNPLA3 was not detected in BAT LDs at 30°C, was detectable only in fasted mice at 22°C, and was present under both conditions at 6°C (Fig. 2B, left). The increase in PNPLA3 was accompanied by a marked decrease in *Pnpla3* mRNA (Fig. 2B, right). Fasting further accentuated the dissociation between *Pnpla3* mRNA and protein levels in BAT (Fig. 2B). Quantitative mass spectrometry confirmed minimal PNPLA3 expression at thermoneutrality, low levels at 22°C (0.3 fmol PNPLA3/μg LD protein), and robust induction at 6°C, especially in fasted mice (Fig. 2C). Cold-induced changes in *Pnpla3* mRNA and PNPLA3 protein levels were comparable in *Pnpla3*^*M/M*^ mice. PNPLA3 protein abundance was higher in *Pnpla3*^*M/M*^ mice than WT mice, as documented previously in liver (Fig. S3A). PNPLA3 levels were also increased in WAT of cold-exposed fasted mice (Fig. S4A).

**Fig. 2.**
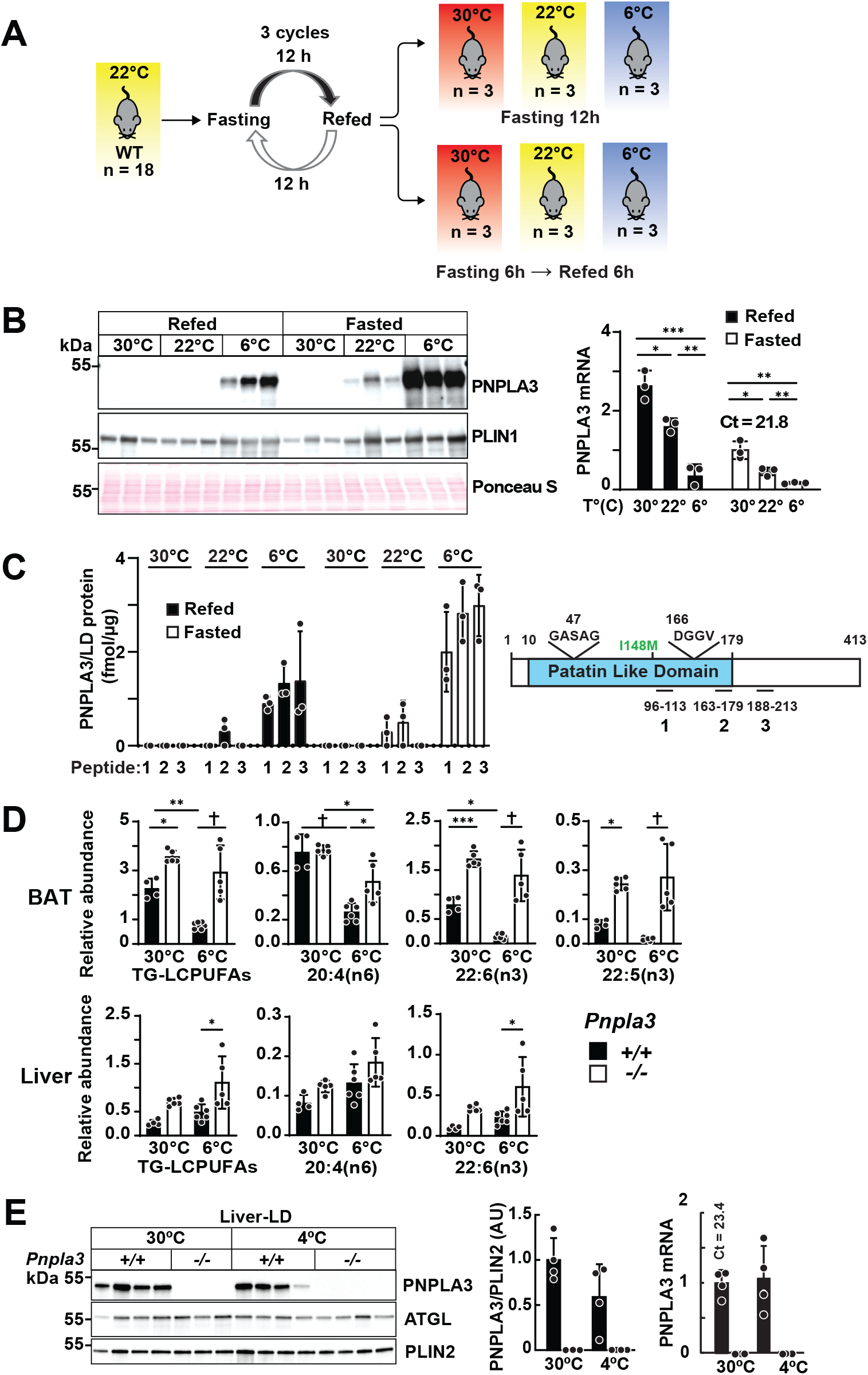
Cold exposure increases PNPLA3 while decreasing *Pnpla3* mRNA levels in BAT. (A) Experimental design. (B) PNPLA3 protein and *Pnpla3* mRNA levels in BAT of WT mice maintained at 6°C, 22°C, or 30°C for 12 h under fasting or refeeding conditions. (C) PNPLA3 quantification in BAT by selected reaction monitoring using three PNPLA3-specific, isotope-labeled peptides. (D) TG-LCPUFA profiles in BAT and liver of WT and *Pnpla3*^*-/-*^ mice (n=4– 6/group) maintained at 30°C or 6°C. (E) PNPLA3 protein in hepatic LDs from mice (n=3–4/group) maintained at 30°C or 4°C. Data represent mean ± SD. Statistical analysis was performed using one-way ANOVA with Tukey’s multiple comparisons test. ^*^P < 0.05; ^**^P < 0.01; ^***^P < 0.001; †P < 0.0001.

### PNPLA3 expression in BAT is associated with a decrease in TG-LCPUFAs and reduced circulating free LCPUFAs

Since PNPLA3 expression in liver is associated with reduced TG-LCPUFAs levels^[4]^, selected TG-LCPUFAs were measured in BAT from WT and *Pnpla3*^*-/-*^ mice housed at 30°C or 6°C for one week. Cold exposure reduced TG-LCPUFAs by ∼69% in WT mice but had no effect in *Pnpla3*^*-/-*^ mice (Fig. 2D, top). No reductions in TG-LCPUFAs or changes in other FAs were observed in livers of cold-exposed mice (Fig. 2D, bottom and Fig. S4B), or in WAT depots of cold-exposed mice (Fig. S4C). The lack of any change in TG-LCPUFAs in WAT in this experiment may be due to lower levels of PNPLA3 expression.

Seven days of cold exposure did not alter circulating free LCPUFA levels in WT mice, but increased plasma LCPUFAs ∼4-fold in *Pnpla3*^*-/-*^ mice (Fig. S4D).

### No change in hepatic *Pnpla3* mRNA or protein levels with cold exposure

To test whether hepatic PNPLA3 is regulated by cold, *Pnpla3* mRNA and LD associated PNPLA3 protein in liver of mice housed at 30°C or 4°C for 12 h were measured. Neither PNPLA3 abundance nor hepatic FA composition changed with cold exposure (Fig. 2E and Fig. 2D, bottom). Thus, cold exposure exerts an adipose tissue-specific effect on PNPLA3 expression.

### Metabolic responses to cold are preserved in *Pnpla3*^*-/-*^ mice

To investigate the role of PNPLA3 in energy homeostasis during cold exposure, WT and *Pnpla3*^*-/-*^ mice were housed in metabolic cages at either 30°C or 6°C. The expected cold-induced increases in food intake, oxygen consumption (VO_2_) and carbon dioxide production (VCO_2_), as well as decreases in respiratory exchange ratio (RER), were observed, with no differences between genotypes (Fig. 3A–D). Rectal temperatures were also similar at both ambient temperatures (Fig. 3E), indicating that PNPLA3 deficiency does not impair energy expenditure or thermogenesis during cold exposure. No significant differences in levels of circulating TGs or free fatty acids were found in WT and *Pnpla3*^−*/*−^ mice under either condition (Fig. S5).

**Fig. 3.**
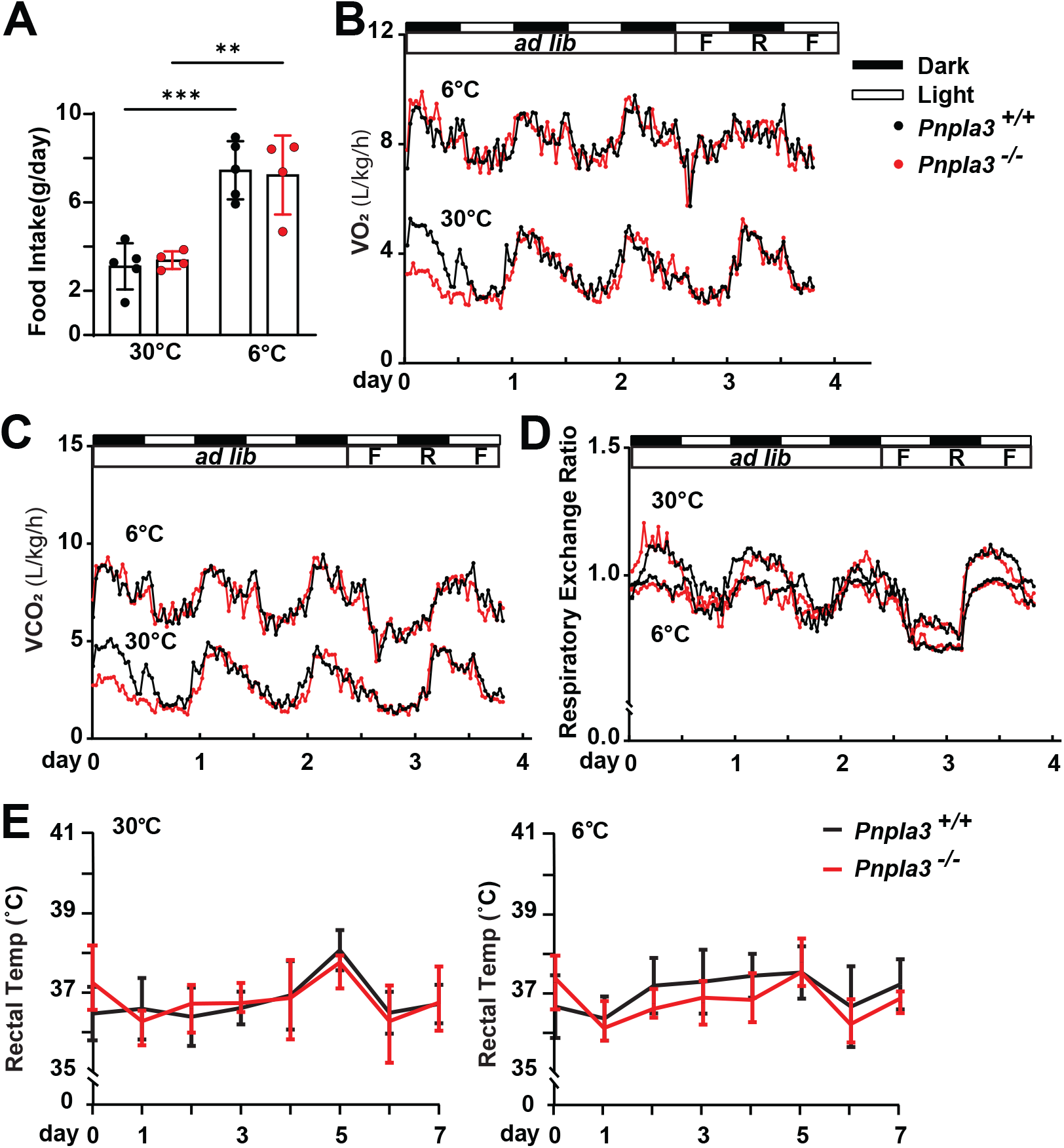
Indirect calorimetry in WT and *Pnpla3*^*-/-*^ mice after cold exposure. (A–D) Food intake (A), VO_2_ (B), VCO_2_ (C), and respiratory exchange ratio (D) in WT and *Pnpla3*^*-/-*^ mice (n=4/group) maintained at 30°C or 6°C. Mice were fed *ad libitum* for 60 h, then subjected to 12-h cycles of food restriction, refeeding, and fasting. (E) Rectal temperature of WT and *Pnpla3*^*-/-*^ mice (n=4–5/group) maintained at 30°C or 6°C. Data represent mean ± SD. Indirect calorimetry data were analyzed using repeated-measures (mixed-effects) ANOVA.

### Activation of the β3-adrenergic receptor (β3AR) increases PNPLA3 in BAT and cultured adipocytes

Because BAT responses to cold are mediated by β3AR signaling^[17]^, mice at 22°C were treated with the selective β3AR agonist CL316243^[18]^. In BAT, PNPLA3 increased 43-fold (Fig. 4A, left), while *Pnpla3* mRNA decreased by 76% (Fig. 4A, right). In contrast, hepatic PNPLA3 levels were unchanged despite a 47% reduction in hepatic *Pnpla3* mRNA. *Atgl* mRNA and ATGL protein were unaffected in both tissues (Fig. 4A, bottom), indicating selective β3AR–dependent regulation of PNPLA3 in BAT. A similar dissociation between the levels of PNPLA3 and *Pnpla3* mRNA was observed with CL316243 treatment in V-WAT: a 1.8-fold increase in PNPLA3 and 84% reduction in *Pnpla3* mRNA levels (Fig. S6A). To further define the temporal relationship between β3AR signaling and PNPLA3 levels, we performed a time-course experiment after giving CL316243. In both BAT and WAT, *Pnpla3* mRNA levels declined within 30 min and continued to decrease over time (Fig. S6B). In BAT, PNPLA3 levels increased at the 30 min timepoint and reached a maximum at 6 h, whereas in WAT the increase in PNPLA3 occurred with slower kinetics but followed a similar trend. In contrast, despite a reduction in *Pnpla3* mRNA levels, hepatic PNPLA3 levels did not change. These findings demonstrate a rapid and tissue-specific dissociation between *Pnpla3* mRNA and protein abundance in response to β3AR stimulation.

**Fig. 4.**
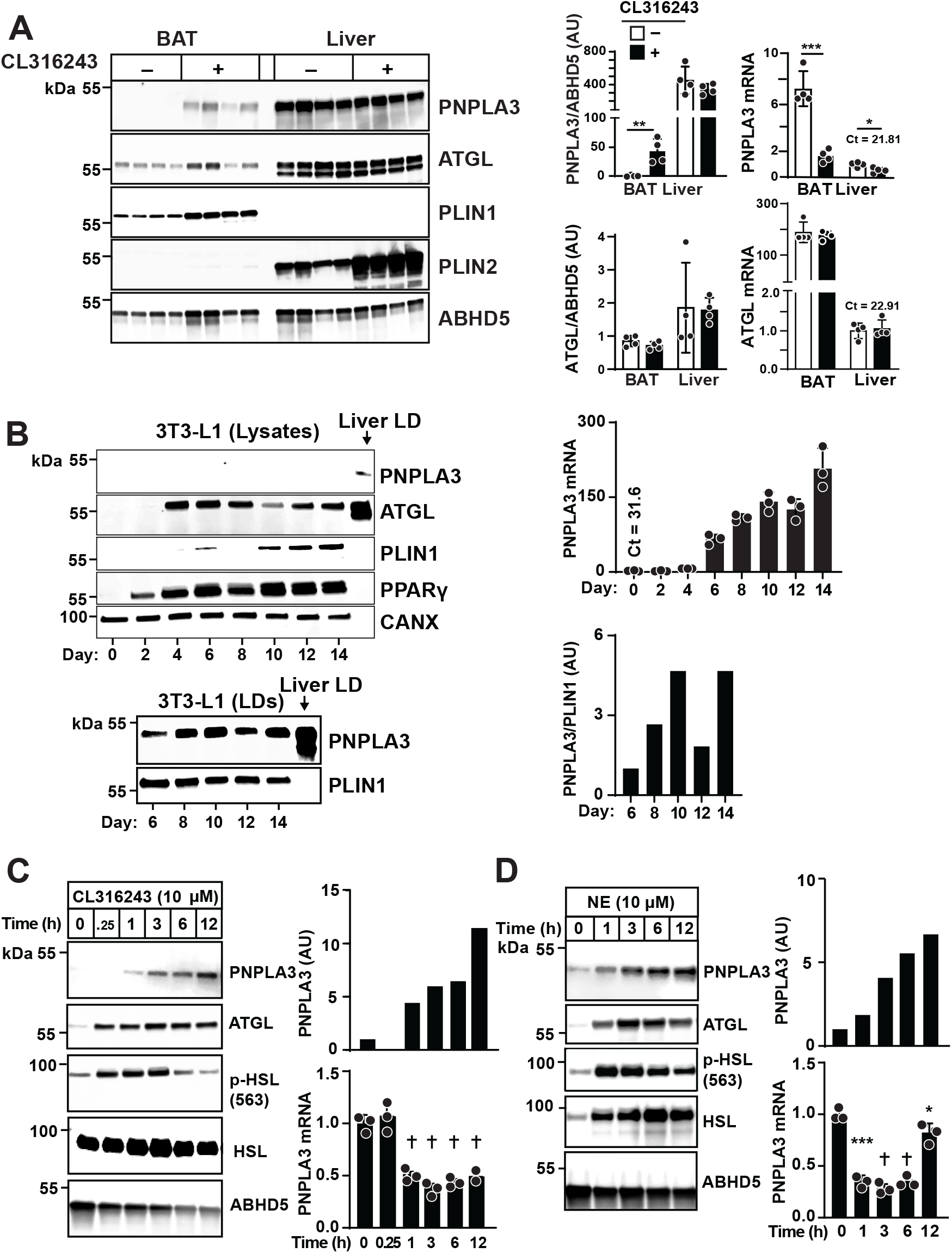
β3AR agonists increase PNPLA3 protein and decrease *Pnpla3* mRNA in BAT and adipocytes. (A) PNPLA3 protein and *Pnpla3* mRNA in BAT and liver from WT mice (n=4/group) treated with CL316243 or saline. (B) PNPLA3 and *Pnpla3* mRNA levels in differentiating 3T3-L1 cells. (C, D) PNPLA3 protein in LDs and *Pnpla3* mRNA levels in mature 3T3-L1 cells treated with CL316243 or norepinephrine (NE). Data represent mean ± SD. Statistical analyses were performed using Student’s *t*-test (A) or one-way ANOVA with Dunnett’s multiple comparisons test (C, D). ^*^P < 0.05; ^**^P < 0.01; ^***^P < 0.001; †P < 0.0001.

To further define the signaling pathway(s) responsible for cold-induced changes in PNPLA3, we established a cell culture model using murine preadipocytes (3T3-L1 cells). During 3T3-L1 differentiation^[19]^, *Pnpla3* mRNA levels increased ∼200-fold while PNPLA3 protein was undetectable using whole-cell lysates (Fig. 4B, top). We repeated the experiment using LDs, which could be isolated as early as day 6, and PNPLA3 was readily detected (Fig. 4B, bottom). Treatment of differentiated 3T3-L1 cells (day 12) with CL316243 increased PNPLA3 levels 4.4-fold at 1 h and 11.5-fold at 12 h, whereas *Pnpla3* mRNA levels fell by 50% (Fig. 4C). Norepinephrine (NE) similarly elevated PNPLA3 levels (6.7-fold at 12 h), while *Pnpla3* mRNA levels fell to 28% of baseline by 3 h (Fig. 4D). Thus, *Pnpla3* mRNA and protein levels were also dissociated following β-adrenergic receptor (βAR) stimulation in cultured adipocytes.

### NE stimulates PNPLA3 via cAMP/PKA and PI3K/AKT signaling pathways

To determine if the NE-stimulated increase in PNPLA3 protein is mediated via the cyclic AMP (cAMP)–protein kinase A (PKA) pathway, we treated 3T3-L1 cells with forskolin, an adenylyl cyclase activator^[20]^. Forskolin rapidly increased PNPLA3 and ATGL levels and stimulated phosphorylation of hormone-sensitive lipase (HSL) (Fig. 5A, left). Coincident with the increase in PNPLA3 protein, levels of *Pnpla3* mRNA fell. The PKA inhibitor H-89^[21]^ blocked this effect (Fig. S7A). Treatment with 8-Bromo-cAMP, a cAMP analog, also increased PNPLA3 levels while decreasing *Pnpla3* mRNA levels (Fig. S7B). NE-induced PNPLA3 upregulation was blocked by inhibitors of phosphatidylinositol 3-kinase (PI3K; LY294002)^[22]^, AK strain transforming 1 kinase (AKT; AKTi VIII)^[23]^, and Torin 1, an inhibitor of mTOR complex 1 (mTORC1) and 2 (mTORC2)^[24]^ (Fig. 5B–D). In contrast, rapamycin, which inhibits only mTORC1^[25]^, did not interfere with the NE-induced increase in PNPLA3 (Fig. S7C). The finding that Torin 1, but not rapamycin, inhibited NE-induced increases in PNPLA3 implicates mTORC2 in contributing to this response.

**Fig. 5.**
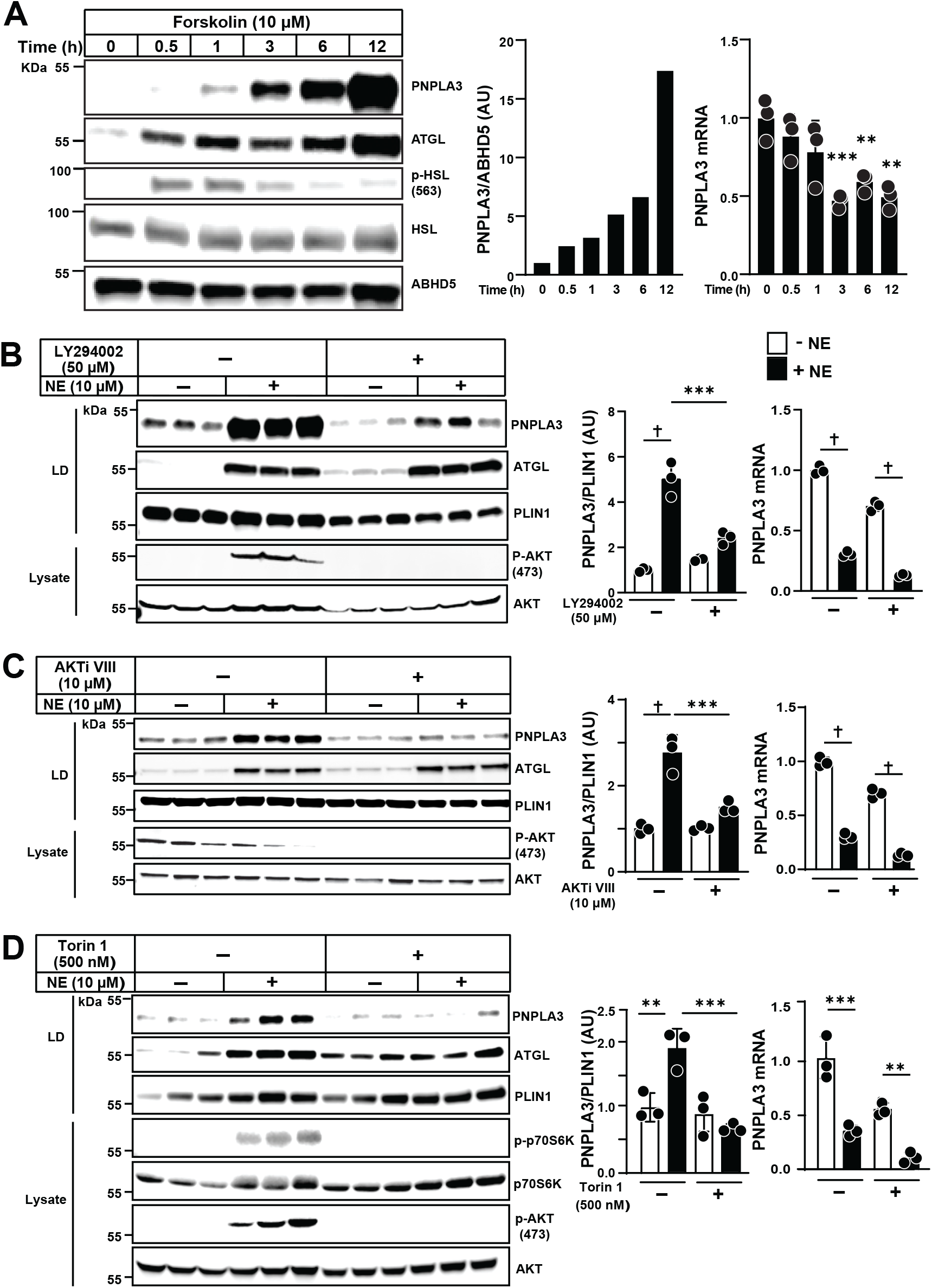
PNPLA3 expression in adipocytes is regulated by cAMP/PKA and PI3K/AKT signaling pathways. (A) PNPLA3 protein on LDs (left and middle) and *Pnpla3* mRNA (right) in 3T3-L1 adipocytes treated with forskolin. (B–D) PNPLA3 protein on LDs (left and middle) and *Pnpla3* mRNA (right) in 3T3-L1 adipocytes treated with norepinephrine (NE) with or without LY294002 (B), AKTi VIII (C), and Torin 1 (D). Data represent mean ± SD. Statistical analyses were performed using one-way ANOVA with Dunnett’s multiple comparisons test (A) and Tukey’s multiple comparisons test (B–D). ^*^P < 0.05; ^**^P < 0.01; ^***^P < 0.001; †P < 0.0001.

### Adrenergic signaling reduces PNPLA3 degradation

To assess whether NE increases PNPLA3 levels by reducing its degradation, we inhibited protein synthesis using cycloheximide (CHX) in 3T3-L1 cells and monitored PNPLA3 levels over the ensuing 100 min^[26]^. Under basal conditions, PNPLA3 levels remained relatively stable over 100 min (Fig. 6A). In the presence of CHX, PNPLA3 levels declined more rapidly (T_1/2_ = 40 min). Co-treatment with NE and CHX extended the half-life of PNPLA3 (T_1/2_ > 100 min). These results are consistent with NE stabilizing PNPLA3 by reducing its degradation, although it remains possible that the effect of CHX was indirect.

**Fig. 6.**
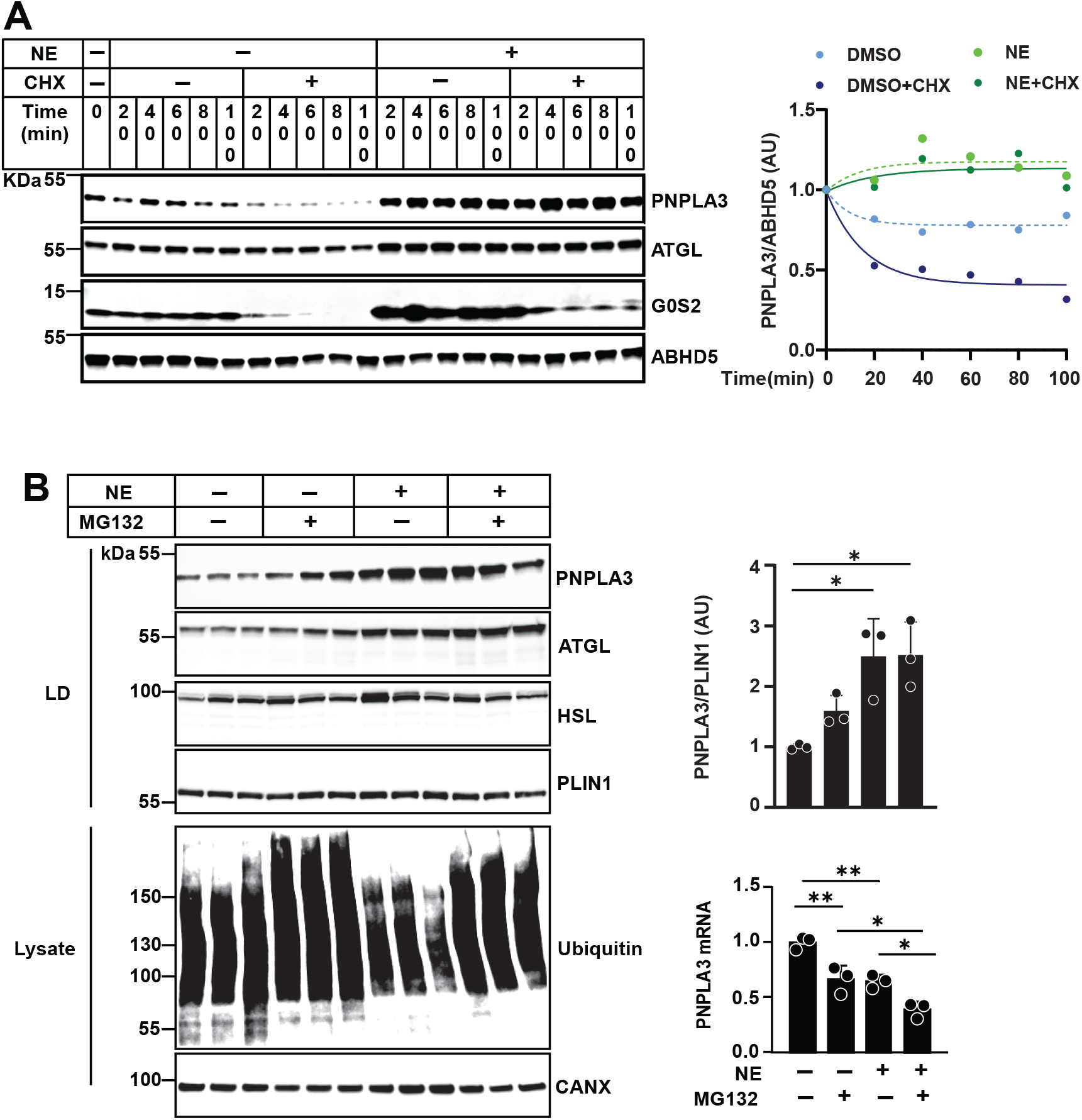
β3AR stimulation increases PNPLA3 protein by inhibiting ubiquitin-proteasome-mediated degradation. (A) PNPLA3 protein on LDs (left) and corresponding degradation curves (right) in 3T3-L1 adipocytes treated with norepinephrine (NE), cycloheximide (CHX), or both. (B) PNPLA3 protein on LDs (left and upper right) and *Pnpla3* mRNA levels (lower right) in 3T3-L1 adipocytes treated with NE with or without MG132. Data represent mean ± SD. Statistical significance was assessed using one-way ANOVA with Tukey’s multiple comparisons test; ^*^P < 0.05; ^**^P < 0.01.

PNPLA3 in liver undergoes ubiquitin-dependent proteasomal degradation^[8, 27]^. Therefore, we treated 3T3-L1 cells with NE in the absence or presence of the proteasomal inhibitor MG132^[28]^. Treatment with MG132 for 3 h resulted in a 1.6-fold increase in PNPLA3 (Fig. 6B). NE further increased PNPLA3 levels, but addition of MG132 to NE-treated cells did not cause any additional increase in PNPLA3. Thus, NE stabilizes PNPLA3, likely by inhibiting its degradation via the ubiquitin-proteasome pathway. We cannot rule out the possibility that enhanced translation also contributes to the NE-induced PNPLA3 accumulation.

### Reduced ribosome association of *Pnpla3* mRNA in BAT of cold-exposed mice

We excluded several mechanisms that might contribute to the divergence between *Pnpla3* mRNA and protein levels associated with cold exposure in BAT, including cold-induced differences in *Pnpla3* mRNA splicing (Fig. S8A), polyadenylation (Fig. S8B), or trafficking out of the nucleus (Fig. S8C). We also tested if cold exposure alters translational efficiency of *Pnpla3* mRNA by promoting its association with ribosomes. We used mice expressing a small ribosomal protein L22 (RPL22) tagged with hemagglutinin (HA) in adipocytes but not in liver (Fig. 7A, 7B)^[29, 30]^. Immunoprecipitation of RPL22-HA recovered intact ribosomes (Fig. 7C). Quantitative RT-qPCR revealed that ∼49% of *Pnpla3* mRNA was bound to ribosomes in mice kept at 30°C, and this fraction decreased by 57% following cold exposure (Fig. 7D), whereas ribosome association of other mRNAs remained unchanged. Therefore, the increase in PNPLA3 in BAT of cold-exposed mice is not a consequence of increased association of *Pnpla3* mRNA with ribosomes.

**Fig. 7.**
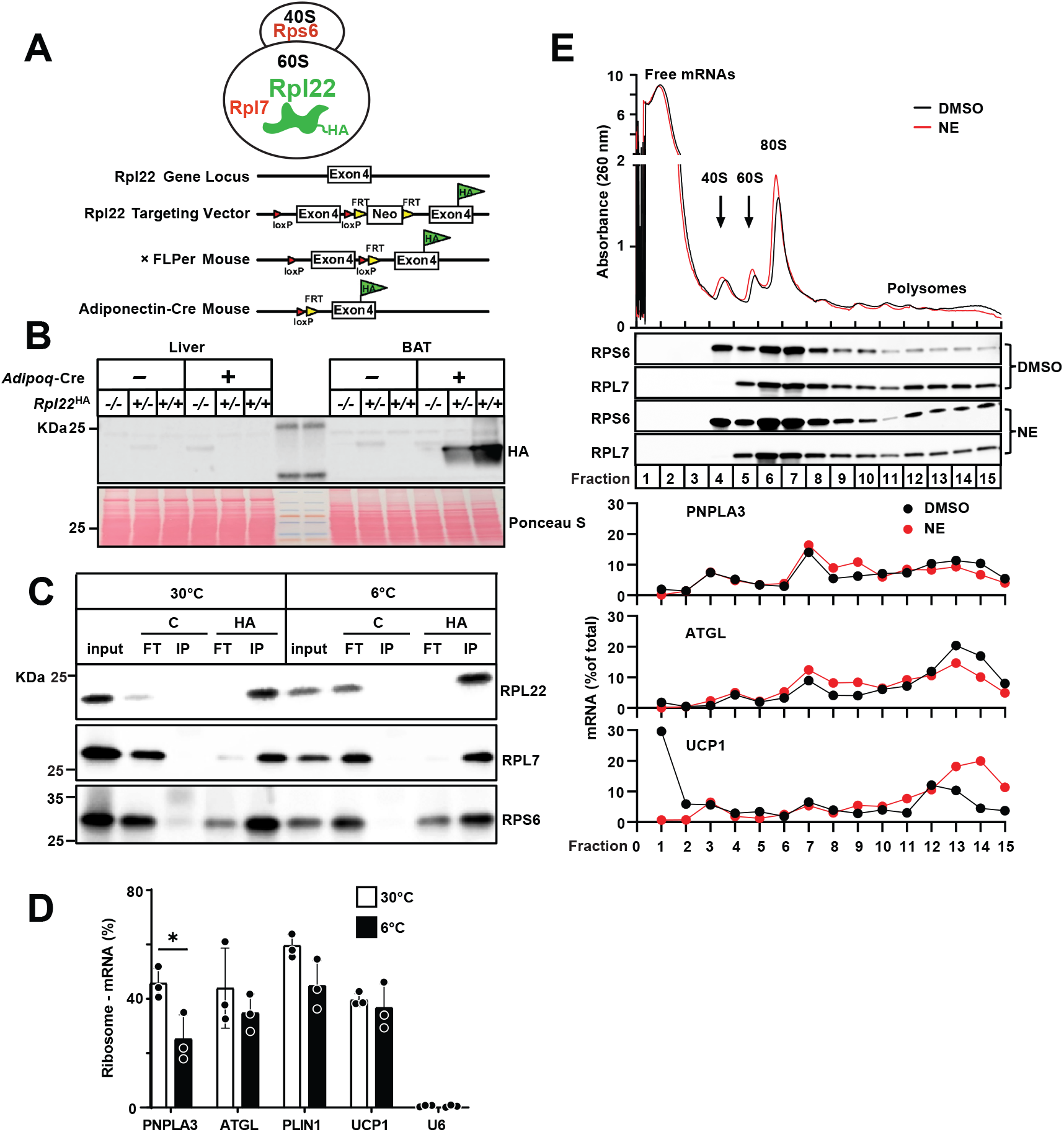
Binding efficiency of *Pnpla3* mRNA to ribosomes in adipose tissue and adipocytes. (A) Generation of adipose-specific RPL22-HA RiboTag mice. (B) RPL22-HA expression in liver and BAT. (C) Detection of ribosomal proteins after HA-based immunoprecipitation. (D) Ribosome-associated mRNA in BAT from mice (n=3/group) maintained at 30°C or 6°C for 12 h. (E) Polysome profiling of 3T3-L1 adipocytes treated with norepinephrine (NE) or DMSO, showing absorbance traces (upper), ribosomal subunit distribution (middle), and distribution of target mRNAs across fractions (lower). Data represent mean ± SD. Statistical analysis was performed using a Student’s *t*-test; ^*^P < 0.05.

To test if NE promoted polysome assembly on *Pnpla3* transcripts to increase translation, we separated ribosome fractions from DMSO- and NE-treated cells by sucrose density gradient centrifugation (Fig. 7E). Absorbance at 260 nm revealed well-resolved peaks (Fig. 7E, top) that did not differ between DMSO- and NE-treated groups, suggesting that overall translational activity was not markedly altered. We validated the identity of ribosomal complexes by subjecting gradient fractions to immunoblotting for ribosomal proteins RPS6 and RPL7, which were present in the polysome fractions. The distribution of *Pnpla3* mRNA across polysome fractions was unchanged by NE treatment, whereas *Ucp1* mRNA polysome association increased, as expected^[31]^. Thus, increased PNPLA3 protein is not due to enhanced ribosome association or polysome loading during adrenergic stimulation.

## Discussion

The major finding of this study is that PNPLA3 regulation differs markedly between liver and adipose tissue in response to distinct stimuli acting through different signaling pathways. It is well established that *Pnpla3* mRNA and protein are expressed at very low levels in the livers of fasting mice and that both increase dramatically upon refeeding via SREBP-1c–mediated transcriptional activation^[7]^. In BAT, *Pnpla3* mRNA changes with feeding are more modest, and PNPLA3 was undetectable at thermoneutrality, despite BAT having the highest *Pnpla3* mRNA abundance (Fig. 1B). Cold exposure did not induce changes in hepatic PNPLA3 (Fig. 2E), yet it caused a 19- and 40-fold increase in PNPLA3 in BAT from fed and fasted mice. The increases in PNPLA3 occurred despite a coordinated >75% reduction in *Pnpla3* mRNA (Fig. 2B). Similar changes were seen in WAT. These findings reveal major tissue-specific differences in regulation of PNPLA3 expression and provide a framework for identifying the physiological role of the enzyme and understanding why the 148M variant confers susceptibility to SLD.

Divergent actions of SREBP-1c on PNPLA3 expression in liver and adipose tissue in response to food intake reflect differences in its function in these two tissues. In liver, PNPLA3 is primarily regulated by SREBP-1c at the transcriptional level, resulting in a marked increase in both *Pnpla3* mRNA and protein levels in response to food intake^[7]^. In contrast, fasting/refeeding only modestly altered *Pnpla3* mRNA levels in adipose tissue (Fig. 1B). These tissue-specific differences in SREBP-1c responsiveness were noted previously for other SREBP-1c target genes^[32]^. SREBP-1c fails to bind the sterol regulatory element/E-box in the fatty acid synthase (FAS) promoter in adipocytes, as it does in hepatocytes^[32]^. Moreover, deletion of SREBP-1c in adipocytes does not alter TG metabolism in that tissue^[32]^. Thus, unlike in liver, SREBP-1c– dependent transactivation does not drive PNPLA3 expression or TG metabolism in adipose tissue.

PNPLA3 expression in liver and adipose tissue in response to cold was also discordant^[33]^. Cold exposure did not alter hepatic PNPLA3 levels (Fig. 2E), yet it induced a marked increase in PNPLA3 in BAT and WAT. To gain insight into the physiological role of cold-induced PNPLA3 upregulation in BAT, we characterized the signaling pathways regulating PNPLA3 using well-characterized agonists and inhibitors of key components of major signal transduction cascades. Our data indicated that the cold-induced increase in PNPLA3 (and reduction in *Pnpla3* mRNA) was mediated by β3AR signaling. Both CL316243 (a β3AR agonist) and NE (a relatively nonselective AR agonist) reproduced the cold-induced changes in *Pnpla3* mRNA and protein levels in mouse BAT (Fig. 4A) and in cultured adipocytes (Fig. 4C and 4D). Additional studies using various modulators of the β3AR signaling pathway were consistent with PNPLA3 being a downstream effector of βAR signaling through both AKT and PKA, perhaps acting through mTORC2, linking cold-induced lipolysis to adaptive lipid remodeling in adipose tissue.

A striking finding of this study was the reciprocal change in *Pnpla3* mRNA and protein levels in adipose tissue under both thermoneutral and cold conditions (Fig. 1A and 2B). The discordant regulation resembles that observed for other stress-responsive proteins where precise control of protein expression is crucial. An example is the hypoxia inducible factor alpha (HIF1A), a transcription factor that plays a key role in response to hypoxia. Levels of *Hif1a* mRNA respond minimally to hypoxia, yet HIF1A protein increases dramatically due to reduced degradation^[34]^. For PNPLA3, the discrepancy between mRNA and protein levels is even more pronounced than observed for HIF1A^[34]^. We propose that acute adrenergic signaling stabilizes PNPLA3 by suppressing ubiquitin–proteasome–mediated degradation, allowing rapid protein accumulation despite declining mRNA levels. In parallel, additional post-transcriptional mechanisms may contribute to the rapid increase in PNPLA3 levels, including sequestration of Pnpla3 transcripts, perhaps in stress granules or other ribonucleoprotein complexes under thermoneutral conditions, followed by their release and translation in response to β-adrenergic stimulation. PNPLA3 abundance is likely controlled by coordinated post-transcriptional and post-translational processes during acute metabolic stress. We found no evidence that increased translational efficiency contributed to cold-induced increases in PNPLA3. Cold did not enhance processes that facilitate mRNA translation, such as mRNA splicing, polyadenylation, or nuclear export (Fig. S8). Ribosome association of *Pnpla3* mRNA decreased, rather than increased, with cold exposure (Fig. 7D), and polyribosome formation was not enhanced for *Pnpla3* mRNA, unlike for *Ucp1* mRNA^[31]^(Fig. 7E).

Two lines of evidence support suppression of PNPLA3 degradation as the major mechanism underlying the cold-induced increase in PNPLA3. First, NE treatment prolonged the half-life of PNPLA3 (from 40 min to >100 min). Second, the increase in PNPLA3 levels in NE-treated adipocytes was not further enhanced by the addition of MG132. Further studies will be required to elucidate the molecular details of how cold exposure interferes with proteasome-mediated PNPLA3 degradation.

A key unanswered question is why PNPLA3 expression is selectively increased in adipose tissue in response to cold. It is noteworthy that the highest frequency of PNPLA3(148M)(∼72%) is found in the Yakut population in northeastern Russia, one of the coldest inhabited regions on Earth ^[35]^. Furthermore, PNPLA3(148M) was fixed in both Neanderthals and Denisovans^[36]^. These hominins inhabited Eurasia during the Ice Age, during prolonged periods of extreme cold and food scarcity. This observation raises the possibility that PNPLA3(148M) may have conferred a selective advantage in conditions of combined thermal and nutritional stress by possibly altering lipid storage and remodeling to promote energy conservation or efficient lipid mobilization. Viewed from this perspective, the same PNPLA3 variant that predisposes individuals to SLD in contemporary obesogenic environments may have served an adaptive role in ancestral settings characterized by cold exposure and limited energy availability.

Generating sufficient heat to maintain body temperature is a major challenge for mice^[37]^, which adapt to cold by uncoupling oxidative phosphorylation^[16]^. Lowering ambient temperature from 30°C to 6°C doubled VO_2_, but the response did not differ between WT and *Pnpla3*^*-/-*^ mice (Fig. 3). Indeed, we failed to identify any defects in cold adaptation in *Pnpla3*^*-/-*^ mice. Importantly, these experiments were conducted under a single set of experimental conditions. Therefore, we cannot exclude the possibility that PNPLA3-dependent phenotypes may emerge under alternative conditions (e.g., different diets or environmental exposures). Further studies will be required to systematically explore these possibilities.

The only difference in phenotype we observed between WT and *Pnpla3*^*-/-*^ mice was an enrichment of TG-LCPUFAs in the tissues of *Pnpla3*^*-/-*^ mice (Fig. 2D). Rather than acting as a direct driver of heat production, PNPLA3 may function primarily as a regulator of lipid remodeling. Consistent with this notion, PNPLA3 expression drives the redistribution of LCPUFAs between TGs and PLs on LDs^[4]^. The redistribution of LCPUFAs may improve access of lipases to TG for lipolysis. Alternatively, changes in lipid composition may stimulate interactions between LDs and other cellular organelles, such as mitochondria.

Further investigation will also be required to determine how PNPLA3(148M) expression in adipose tissue alters lipid homeostasis. Expression of PNPLA3(148M) promotes lipid remodeling in both humans and mice, though the patterns of changes in TG-LCPUFAs enrichment differ between the species^[10, 38]^. In humans, PNPLA3(148M) is associated with an increase in TG-LCPUFAs, which is similar to what is seen in *Pnpla3*^*-/-*^ mice^[10, 38]^. Conversely, hepatic and adipose tissue TGs of mice expressing PNPLA3(148M) are depleted in LCPUFAs. These species-specific differences may reflect divergent effects of the 148M variant on PNPLA3 function in mice versus humans. In biochemical studies using purified enzyme, the 148M variant causes a ∼80% reduction in the TG hydrolase activity. In mice, PNPLA3(148M) accumulates to ∼40-fold higher levels than the WT protein, which may compensate for the reduction in lipase activity^[14]^. If PNPLA3(148M) does not accumulate to similarly high levels in human tissues, then the lipid pattern would be expected to resemble that seen in *Pnpla3*^*-/-*^ mice, which is what we observed. Additional studies will be required to elucidate how the changes in lipid composition of both hepatic and adipose tissue LDs relate to the increased risk of SLD associated with PNPLA3(148M).

## Abbreviations

The abbreviations are detailed in the *Supplementary Data*.

## Data availability statement

All data presented in this manuscript will be made available upon request.

## Acknowledgements

We thank Lisa Kinch, Julia Kozlitina, and Benjamin Tu for helpful discussions, and Christina Zhao, Fang Xu and Tommy Hyatt for excellent technical support. We also thank Jeffrey McDonald, Goncalo Dias do Vale, Ann Johnson, Andrew Lemoff and Xuemei Luo for lipid and protein measurements.

## Declaration of generative AI and AI-assisted technologies in the manuscript preparation process

To meet formatting and word-count requirements of the *Journal of Hepatology*, we condensed the text relative to the bioRxiv preprint and in this process used generative AI (Perplexity). After using this tool, the authors reviewed and edited the content as needed and took full responsibility for the content of the published article.

## Authors’ contributions

P.W., Y.W., J.C.C., and H.H.H. designed the experiments; P.W. and Y.W. conducted the experiments; P.W., Y.W., J.C.C. and H.H.H. analyzed the data and wrote the paper.

