## supplementary data for "Tissue-specific regulation of PNPLA3 promotes lipid remodeling in response to dietary and environmental challenges"

**Table of contents**

Materials and Methods……………………………………………………………………..2

Abbreviations………………………………………………………………………………...13

Supplementary Figures……………………………………………………………………15

Supplementary Tables………………………………………………………………………...29

Supplementary References………………………………………………………………....37

### Materials and Methods

#### *Materials*

Please see Table S1.

#### *Mice*

*Pnpla3* I148M (*Pnpla3$^{M/M}$*) and *Pnpla3* S47A (*Pnpla3$^{A/A}$*) knock-in mice were generated as previously described[1]. *Pnpla3$^{-/-}$* mice were a generous gift from Erin Kershaw (University of Pittsburgh, Pittsburgh, PA)[2]. RiboTag mice (*Rpl22-HA$^{fl/fl}$*; strain #: 029977) were obtained from The Jackson Laboratory and crossed with mice expressing Cre under the control of the adiponectin promoter (*Adipoq*-Cre; strain#: 028020) to obtain $F_1$ *Rpl22-HA$^{fl/+}$* Tg(*Adipoq*-Cre) offspring[3]. $F_1$ mice were then crossed with *Rpl22-HA$^{fl/+}$* mice to generate *Rpl22-HA$^{fl/fl}$* Tg(*Adipoq*-Cre) mice. Genotypes were confirmed by PCR-based genotyping using flanking oligonucleotides (available upon request).

All mice were bred onto a C57BL/6J background for more than 3 generations and housed in a standard animal facility under a 12 h light/12 h dark cycle (lights on 7:00 a.m.–7:00 p.m.). Mice were fed a chow diet (PicoLab Rodent Diet 5053) *ad libitum* and maintained at 20–22°C. For dietary challenge experiments, mice were fed a high-sucrose diet (HSD; Fisher Scientific/MP Biomedicals™ Fat Free Diet) for 4 weeks. Prior to experiments, mice underwent dietary synchronization for 3 days consisting of 12 h fasting followed by 12 h refeeding. Unless otherwise specified, mice were euthanized after the final refeeding cycle. For temperature challenge studies, mice were housed individually in incubators maintained at either 4–6°C or 28–30°C, with free access to water and food as specified by the experimental design.

For CL316243 treatment, mice were housed at room temperature (20–22°C) and maintained on a HSD. After 3 days of dietary synchronization, mice received an intraperitoneal injection of CL316243 (1 mg/kg) at the end of the final fasting cycle, after which food was provided. Mice were euthanized 6 h later, and liver and brown adipose tissue (BAT) were collected and immediately placed in liquid $N_2$.

A combined indirect calorimetry system (CaloSys Calorimetry System; TSE Systems, Inc., Bad Homburg, Germany) was used for all metabolic studies. Mice were acclimated to individual metabolic cages for 3 days before data acquisition. For measurements, cages were placed in temperature-controlled incubators set to 6°C, 22°C, or 30°C. Animals were monitored continuously for 96 h, consisting of 60 h with *ad libitum* access to a HSD, followed by a 12 h fasting period, a 12 h refeeding period with HSD, and a final 12 h fasting period. Oxygen consumption ($VO_2$), carbon dioxide production ($VCO_2$), and food intake were recorded continuously, and metabolic rate was normalized to body weight.

All animal experiments were conducted in accordance with protocols approved by the Institutional Animal Care and Use Committee of the University of Texas Southwestern Medical Center.

*Cell culture*

3T3-L1 murine fibroblasts were grown to confluence in growth medium [high glucose DMEM supplemented with 10% fetal calf serum (FCS), penicillin (100 units/mL), and streptomycin (100 µg/mL)] at 37°C in 8.8% $CO_2$. Two days after reaching confluence (designated as day 0), differentiation was induced by adding insulin (5 µg/mL), dexamethasone (1 µM), 3-isobutyl-1-methylxanthine (IBMX, 0.5 mM), and rosiglitazone (1 µM) and cells were grown for an additional 48 h[4]. Subsequently, cells were refed every other day with growth medium containing insulin

(5 μg/mL). Full differentiation into mature adipocytes typically occurred within 10–12 days. Insulin was withdrawn from mature adipocytes for two days prior to the start of the experiment, unless otherwise stated. The 3T3-L1 cell line used in these experiments expresses PNPLA3(WT).

Cells were treated with a single dose of norepinephrine (NE, 10 μM), CL316243 (10 μM), forskolin (10 μM), or 8-Bromo-cAMP (1 mM) in high glucose DMEM supplemented with 2% fatty acid–free bovine serum albumin (FAF-BSA) and harvested at the indicated time points.

For PKA inhibition experiments, cells were pretreated with H-89 (50 μM) for 1 h, followed by treatment with forskolin (5 μM) or vehicle (DMSO) in the presence or absence of H-89, using fresh medium containing 2% FAF-BSA. After 3 h of incubation, cells were harvested for analysis.

For kinase inhibitor experiments, cells were incubated overnight with LY294002 (50 μM), AKTi VIII (10 μM), Torin 1 (500 nM) or rapamycin (100 nM). The following day, cells were stimulated with NE (10 μM) or vehicle (DMSO) in fresh medium containing 2% FAF–BSA, in the presence or absence of the inhibitor. After 3 h, cells were harvested for analysis.

For proteasome inhibition experiments, cells were treated with NE (10 μM) or DMSO in the absence or presence of MG132 (10 μM) in medium containing 2% FAF–BSA for 3 h. For protein synthesis inhibition, cells were treated with cycloheximide (CHX, 10 μM) together with NE or DMSO on day 12 post-differentiation without prior insulin starvation.

*Lipid droplet isolation*

Lipid droplets (LDs) from BAT were isolated as previously described[5] with slight modifications. BAT was homogenized in 1.5 mL ice-cold Buffer A [25 mM Tricine (pH 7.6), 250 mM sucrose, and protease inhibitors and passed through a 200-μm mesh. The homogenates were centrifuged at 2,000 *g* for 15 min at 4°C to isolate LDs. The remaining homogenates were centrifuged at 12,000 *g* for 15 min to obtain smaller LDs. The LDs were further purified by centrifugation at the same respective speeds (2,000 *g* or 12,000 *g*) for 3 min, then resuspended in 200 µL of Buffer B [20 mM HEPES (pH 7.4), 100 mM KCl, 2 mM $MgCl_2$, and protease inhibitors] and vortexed. Washing was repeated twice, and LD proteins were precipitated by adding cold acetone followed by centrifugation at 15,000 *g* for 15 min. The protein pellet was washed with acetone/ethyl ether (50:50, v/v) solution and air-dried. The dried pellet was dissolved in PBS containing 1 M urea and 2% (v/v) SDS. After solubilization, BAT LD proteins from the same mice were pooled[6]. The same method was used to isolate fat cake from subcutaneous white adipose tissue (SQ-WAT) and visceral white adipose tissue (V-WAT). Hepatic LDs were isolated and purified as previously described[6].

Adipocyte LDs were isolated as previously described with slight modifications[7]. Washed adipocytes were incubated with ice-cold Buffer A and Benzonase® Nuclease. Cells were disrupted by repeated pipetting and then placed on ice for 10 min. The suspended cells were transferred to a Dounce homogenizer and gently homogenized on ice using 10 strokes of a hand-held pestle. The homogenate was transferred to an Eppendorf tube and centrifuged at 1,000 *g* for 10 min at 4°C. The supernatant and floating fat layer were collected into separate tubes. One-third volume of ice-cold Buffer A containing 60% sucrose (final concentration: 20%) was added to the supernatant and gently mixed. An ultracentrifuge tube (Beckman #344059) was layered with 4.5 mL of Buffer B, followed by 5.5 ml of Buffer A carefully introduced beneath Buffer B. The cell lysate was then added to the bottom of the tube. The sample was centrifuged at 20,000 *g* for 30 min at 4°C. LDs were collected and washed three times with Buffer B. LD proteins were precipitated and solubilized using the same method described above.

*Recombinant adenoviruses generation and delivery in mice*

Adenoviruses expressing either an empty vector or a fusion protein of PNPLA3 with C-terminal V5 and His tags were constructed as previously described[6]. A total of $1.5 \times 10^{11}$ recombinant adenoviral particles in 200 µl of saline were administered to mice via tail vein injection. Following infection, mice were subjected to three cycles of dietary synchronization and killed after the final feeding cycle.

*RNA extraction and quantitative reverse transcription polymerase chain reaction (RT-qPCR)*

Total RNA was extracted from cells or tissues using an RNA extraction kit (RNeasy Plus Universal Mini Kit) according to the manufacturer's instructions. RNA concentration and purity were determined using a NanoDrop spectrophotometer, and RNA integrity was assessed by agarose gel electrophoresis. For each sample, 1–2 µg of total RNA was reverse transcribed using random hexamer primers and reverse transcription reagents (see Table S1). RT-qPCR was performed using an Applied Biosystems real-time PCR system with SYBR Green detection (see Table S1). Each reaction was carried out in a final volume of 20 µl containing 20 ng of cDNA, 167 nM of each primer, and 10 µl of SYBR Green PCR Master Mix. Thermal cycling conditions were as follows: initial denaturation at 95°C for 10 min, followed by 40 cycles of denaturation at 95°C for 15 s, and annealing/extension at 60°C for 1 min. RNA levels were normalized to *cyclophilin B* or *HPRT* mRNA levels, and relative expression was calculated using the $\Delta\Delta Cq$ method.

*Immunoblotting and protein quantification*

Cells or tissue samples were lysed in RIPA buffer [25 mM Tris-HCl (pH 7.6), 150 mM NaCl, 1% Nonidet P-40, 1% sodium deoxycholate, and 0.1% SDS], and protein concentrations were determined using the Pierce™ BCA Protein Assay Kit. Samples were denatured by heating in 1× Laemmli sample buffer at 95°C for 5 min. Proteins were separated by SDS-PAGE and transferred onto nitrocellulose membranes. Membranes were incubated with primary and secondary antibodies (see Table S1), and signals were detected using SuperSignal™ West Pico PLUS or SuperSignal™ West Femto Maximum Sensitivity Substrate and imaged on Odyssey FC Imager (LI-COR). Band intensities were quantified using Image Studio Lite v5.2. A rabbit anti-mouse PNPLA3 mAb (19A6) was developed against a peptide corresponding to residues 152-309 of PNPLA3[1].

*Measurement of plasma triglyceride and free fatty acids*

Plasma triglyceride and free fatty acid levels were measured using a Glycerol Assay Kit (Millipore Sigma, MAK117) and a Free Fatty Acid Assay Kit (LSBio, LS-K170-100), respectively, according to the manufacturers' instructions. Each sample was analyzed in triplicate.

*Triglyceride fatty acid (TG-FA) analysis by GC-MS*

Approximately 50 mg of tissue was homogenized in methanol/dichloromethane (1:2, v/v) and subjected to three-phase liquid extraction as previously described[8]. Briefly, phase separation was achieved by sequential addition of water, methyl acetate, hexane, and acetonitrile (ACN), followed by vortexing and centrifugation. The upper hexane layer (~98% TG) was collected, dried under nitrogen, and hydrolyzed in methanolic KOH containing deuterated FA standards: $^{2}H_{31}$-16:0, $^{2}H_{8}$-20:4, and $^{2}H_{5}$-22:6n3. Following extraction and derivatization with pentafluorobenzyl bromide and triethylamine in acetone, FAs were analyzed by GC-MS (Agilent 7890/5975C) using electron capture negative ionization in selected ion-monitoring mode.

Peak areas of FAs were normalized to the corresponding internal standards based on chain length: ≤C18 to $^{2}H_{31}$-16:0, C20 to $^{2}H_{8}$-20:4, and C22 to $^{2}H_{5}$-22:6n3). Data processing was performed using Mass Hunter software (Agilent).

*Quantification of PNPLA3 by Selected Reaction Monitoring (SRM)*

Proteins were separated by SDS-PAGE on 4–15% gradient precast gels (Bio-Rad) and visualized by Coomassie Brilliant Blue staining. A ~10 mm gel slice (35–55 kDa) corresponding to the expected molecular weight was excised for SRM analysis. Proteins within the slice were reduced with 20 mM dithiothreitol (DTT), alkylated with 27.5 mM iodoacetamide, and digested overnight at 37°C with trypsin. Peptides were extracted from the gel and dried. The samples were reconstituted and spiked with 100 fmol of each heavy-isotope labeled peptide, corresponding to three tryptic peptides: DGLQESLPDNVHQVISGK (aa 96–113), YVDGGVSDNVPVLDAK (aa 163–179), and STNFFHVNITNLSLR (aa 188–213). Peptides were selected based on their specificity and digestion efficiency. Stable isotope-labeled standards ($^{13}C_6$,$^{15}N_2$-lysine or $^{13}C_6$,$^{15}N_4$-arginine at the C-terminus) were synthesized by 21st Century Biochemicals at >97% purity, as determined by high-performance liquid chromatography (HPLC). Peptides were desalted using an Oasis HLB μElution plate, dried, and reconstituted in 10 μL of 2% ACN/0.1% trifluoroacetic acid in water for SRM analysis.

SRM analysis was performed on an AB Sciex 6500 QTRAP mass spectrometer interfaced with a Thermo Fisher Ultimate 3000 RSLCnano HPLC system. Spiked samples were separated on a Dionex Acclaim PepMap100 C18 reverse-phase column (75 μm × 15 cm) using the Ultimate 3000 RSLCnano HPLC system. The system was controlled by Chromeleon Xpress software (v6.8 SR10) in conjunction with Dionex Chromatography MS Link (v2.12). Peptides were separated at a flow rate of 200 nL/min using the following gradient: 0–25% B (15 min), 25–35% B (5 min), and 35–80% B (5 min). Mobile phase A consisted of 2% ACN and 0.1% formic acid in water, and mobile phase B contained 80% ACN, 10% trifluoroethanol (TFE), 10% water, and 0.1% formic acid.

Mass spectrometric analysis was carried out in positive-ion low-mass mode using a NanoSpray III source equipped with a New Objective precut 360 µ PicoTip emitter (FS360-20-10-N20-10.5CT). The source settings were as follows: curtain gas = 30, ion spray voltage = 2,450 V, ion source gas 1 = 6. Analyst Software v.1.6 was used to run the mass spectrometer, and SRM data were analyzed using Skyline v4.1.

*RNA-seq*

Total RNA was isolated using the RNeasy Plus Universal Mini Kit (Qiagen) and treated with DNase. RNA quality (RIN ≥8.5) and concentration were assessed using an Agilent TapeStation 4200 and a Qubit® 4.0 Fluorometer. For library preparation, 1 µg of RNA was used with the TruSeq Stranded Total RNA LT Kit (Illumina), which includes rRNA depletion, fragmentation, cDNA synthesis, adapter ligation, and PCR amplification according to manufacturer's instructions. Sequencing was performed on illumina NextSeq 2000 platform, generating 25–35 million reads per sample. FASTQ files were quality-checked using FastQC and FastQ_screen, mapped to the hg19 reference genome with Tophat, and duplicate reads were marked using Picard tools. Differential expression analysis was conducted with edgeR (FDR <0.05; fold change >1.5), and gene ontology enrichment was analyzed using DAVID (v6.8).

*Immunoprecipitation of polysomes*

The immunoprecipitation of polysomes was performed as described previously[3]. Briefly, 100 µL of protein G magnetic beads (Dynabeads; Invitrogen) were coupled directly to 10 μL of mouse monoclonal anti-HA antibody (HA.11, ascites fluid; Covance) for 45 min in citrate–phosphate buffer (24 mM citric acid, 52 mM dibasic sodium phosphate, pH 5.0). Antibody-coupled beads were washed once in citrate–phosphate buffer (pH 5.0), and twice in immunoprecipitation buffer [50 mM Tris (pH 7.5), 100 mM KCl, 12 mM $MgCl_2$, 1% Nonidet P-40] before being added to homogenates. BAT was weighed and homogenized using a Dounce homogenizer in polysome buffer [50 mM Tris (pH 7.5), 100 mM KCl, 12 mM $MgCl_2$, 1% Nonidet P-40, 1 mM DTT, 200 U/mL RNasin, 100 μg/mL CHX, and protease inhibitor]. Samples were centrifuged at 10,000 *g* for 10 min to obtain a post-mitochondrial supernatant. Supernatants (400 μL) were added directly to the antibody-coupled protein G magnetic beads and rotated overnight at 4°C. The following day, samples were placed on a magnetic rack on ice and supernatants were recovered before washing the pellets three times for 5 min each in high-salt buffer [50 mM Tris (pH 7.5), 300 mM KCl, 12 mM $MgCl_2$, 1% Nonidet P-40, 1 mM DTT, and 100 μg/mL CHX]. After washing, pellets were saved for immunoblotting and RNA extraction.

To prepare total RNA, 2.5 or 5 volumes of Qiagen RLT buffer were added to the immunoprecipitated pellets or input samples, respectively. Total RNA was isolated according to the manufacturer's instructions using the RNeasy Mini kit (Qiagen) and quantified using a NanoDrop 1000 spectrophotometer (Thermo Scientific). RNA quality was assessed by electrophoresis on 2% agarose gels followed by ethidium bromide staining.

*Polysome profiling and RNA distribution analysis*

Polysome profiling was performed as previously described[9] with minor modifications. Briefly, cells were incubated with 100 μg/mL CHX for 10 min at 37°C to arrest ribosomes on mRNAs.

Cells were then washed with ice-cold PBS containing CHX and collected by centrifugation at 500 *g* for 5 min at 4°C. Cell pellets were lysed using a Dounce homogenizer in ice-cold polysome extraction buffer [20 mM Tris (pH 7.5), 100 mM KCl, 5 mM $MgCl_2$, 0.5% Nonidet P-40, 100 μg/mL CHX, 1:1000 RiboLock RNase inhibitor, and protease inhibitor]. Lysates were incubated on ice for 10 min and then cleared by centrifugation at 12,000 *g* for 10 min at 4°C. The upper lipid layer was carefully removed, and the resulting cytoplasmic supernatant was used for gradient loading. Equal volumes of cytoplasmic lysate (1 mL containing 2 mg total protein) were layered onto pre-formed 10–50% (w/v) sucrose gradients prepared in gradient buffer (20 mM Tris (pH 7.5), 100 mM NaCl, 5 mM $MgCl_2$) using a gradient mixer. Gradients were centrifuged at 190,000 *g* (~39,000 rpm) for 90 min at 4°C in a Beckman SW41Ti rotor. Following centrifugation, sucrose gradients were fractionated at a flow rate of 1 mL/min into 15 fractions with continuous monitoring at 260 nm using a BioComp gradient fractionation system. To assess gradient integrity and ribosomal subunit distribution, proteins from each fraction were analyzed by SDS–PAGE followed by immunoblotting for RPL7 and RPS6. RNA was extracted from each sucrose fraction using TRIzol™ reagent and further purified with the RNeasy Mini Kit (Qiagen) according to the manufacturer's instructions. Purified RNA was reverse transcribed using SuperScript™ IV Reverse Transcriptase, and target mRNA distribution across the gradient was quantified by RT-qPCR using gene-specific primers. The relative abundance of each mRNA was calculated as the percentage of total signal across all 15 fractions.

*Subcellular localization of mRNA in nuclear and cytoplasmic fractions*

Cells were collected and centrifuged at 2,000 *g* for 5 min at 4°C. The resulting cell pellet was resuspended in 300 μL of ice-cold Cell Fractionation Buffer (PARIS™ Kit; Thermo Fisher

Scientific, AM1921) and incubated on ice for 10 min with occasional gentle mixing. Samples were then centrifuged at 500 *g* for 5 min at 4 °C to remove the lipid layer. The remaining lysate and pellet (~300 μL) were combined and thoroughly mixed. To separate cytoplasmic and nuclear fractions, samples were centrifuged at 1,000 *g* for 5 min at 4 °C. The supernatant, containing the cytoplasmic fraction, was collected. The nuclear pellet was washed by resuspending in 100 μL of Cell Fractionation Buffer, followed by centrifugation at 1,000 *g* for 3 min at 4 °C; the supernatant was discarded. RNA was extracted from both fractions, and cDNA synthesis was performed as described previously. RT-qPCR was conducted to quantify mRNA levels. The relative abundance of each mRNA in the nuclear and cytoplasmic fractions was calculated as the percentage of the total signal across both compartments.

*Poly(A) tail length analysis*

The poly(A) tail length of *Pnpla3* mRNA in BAT was determined using the USB® Poly(A) Tail-Length Assay Kit (Affymetrix, 76455) according to the manufacturer's instructions, with minor modifications. Total RNA was extracted from BAT of mice maintained at either 30°C or 6°C using the RNeasy Mini Kit. For each sample, 2 μg of total RNA was subjected to guanosine/inosine (G/I) tailing in a 20 μL reaction at 37°C for 60 min, followed by termination with 2 μL of 10 × Tail Stop Solution. The tailed RNA was reverse transcribed at 44°C for 60 min using the kit-supplied RT primer and reverse transcriptase, followed by heat inactivation at 92°C for 10 min. PCR amplification was performed using a gene-specific forward primer located upstream of the *Pnpla3* polyadenylation site (5′-GCAGAAGGATTGAATGGATACA-3′), paired with either the gene-specific reverse primer (5′-TTTATTATGGACCCTTTTCCCTTA-3′) to generate products terminating before the poly(A) tail, or the universal reverse primer provided in the kit to generate products containing the poly(A) tail. Amplification products were resolved on a 5% polyacrylamide gel in 1× TBE buffer and visualized by ethidium bromide staining under UV illumination. The poly(A) tail length was calculated as the size of the poly(A) PCR product minus the distance between the gene-specific forward primer and the putative polyadenylation site.

*Quantification and statistical analysis*

Data are presented as mean ± SD unless otherwise specified. Statistical comparisons between two groups were performed using a unpaired Students'*t*-test. For multiple group comparisons, one-way ANOVA followed by Dunnett's multiple comparison or Tukey's multiple comparisons test was applied as appropriate. Statistical significance was defined as a $p < 0.05$. Details for statistical analyses performed for each experiment are provided in the figure legends.

All experiments were independently repeated at least three times with comparable results.

**Abbreviations**

AKT, AK strain transforming 1 kinase; ATGL, adipose triglyceride lipase; BAT, brown adipose tissue; βAR, β-adrenergic receptor; β3AR, β3-adrenergic receptor; cAMP, cyclic AMP; CHX, cycloheximide; FA, fatty acid; FAS, fatty acid synthase; HIF1A, hypoxia inducible factor alpha; HSD, high-sucrose diet; HSL, hormone-sensitive lipase; LCPUFAs, long-chain polyunsaturated fatty acids; LD, lipid droplet; mTORC1, mTOR complex 1; mTORC2, mTOR complex 2; NE, norepinephrine; PI3K, phosphatidylinositol 3-kinase; PKA, protein kinase A; PL, phospholipid; PNPLA3, patatin-like phospholipase domain-containing protein 3; RER, respiratory exchange ratio; RPL22, ribosomal protein L22; SLD, steatotic liver disease; SQ-WAT, subcutaneous white adipose tissue; SREBP-1c, sterol regulatory element-binding protein-1c; TG, triglyceride; V-WAT, visceral white adipose tissue; WT, wild-type.

**Supplementary Figures**

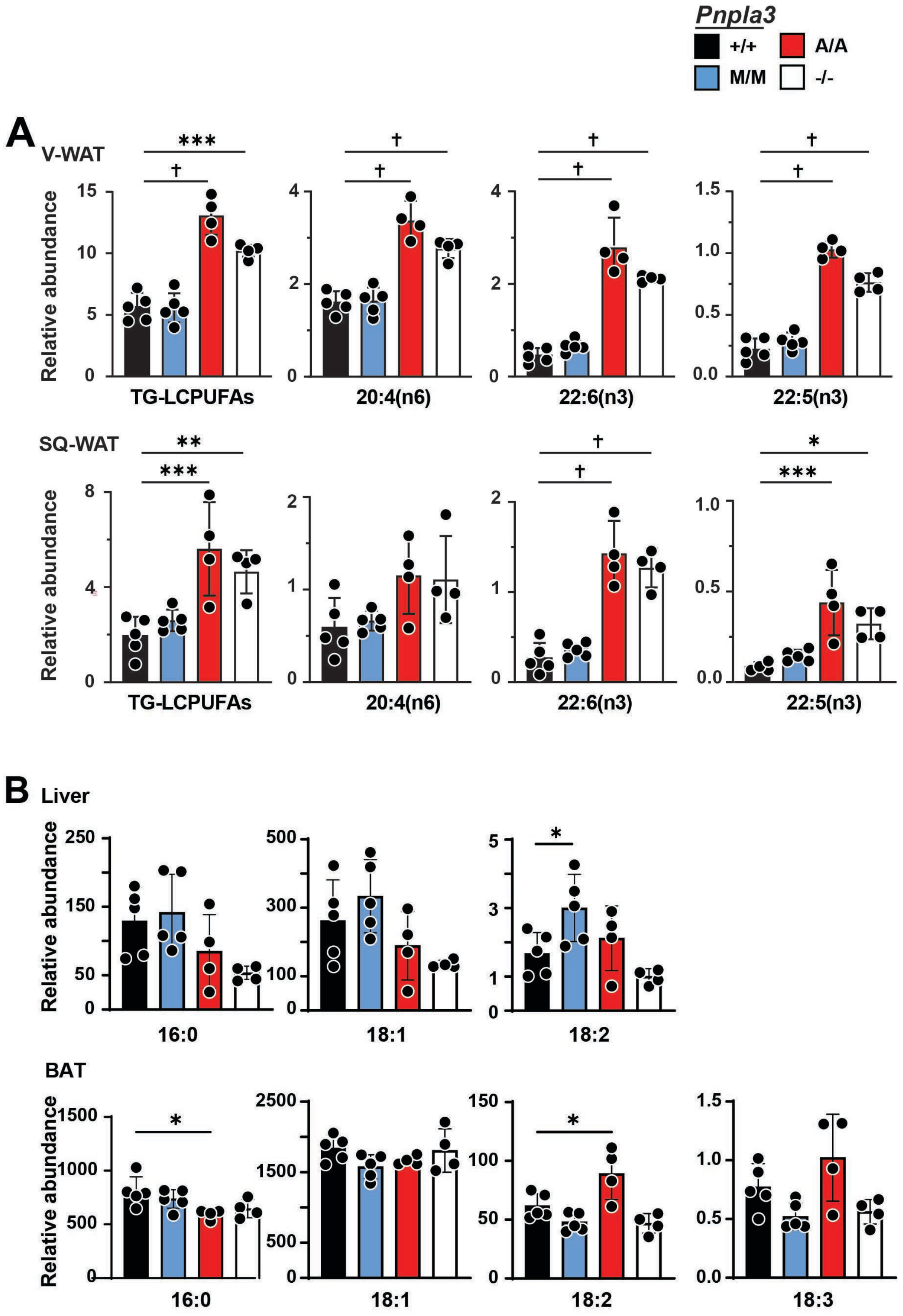


**Fig. S1. Composition of fatty acids (FAs) in triglyceride (TG) from adipose tissue and liver of genetically modified mice.** (A) Relative abundance of TG-long-chain polyunsaturated fatty acids (TG-LCPUFAs) in visceral white adipose tissue (V-WAT, upper) and subcutaneous white adipose tissue (SQ-WAT, lower) from *Pnpla3*$^{-/-}$, *Pnpla3*$^{M/M}$, *Pnpla3*$^{A/A}$ and WT mice. (B) Relative abundance of other FAs (16:0, 18:1, 18:2, 18:3) in TG from liver (upper) and brown adipose tissue (BAT, lower) of the same mice. Mice (n = 4-5/group) were kept at thermoneutrality (30ºC) and fed a high sucrose diet (HSD) for 4 weeks. Data represent mean ± SD. P values were determined by one-way ANOVA with Tukey's multiple comparisons test. * P < 0.05; ** P < 0.01; *** P < 0.001; †P < 0.0001.

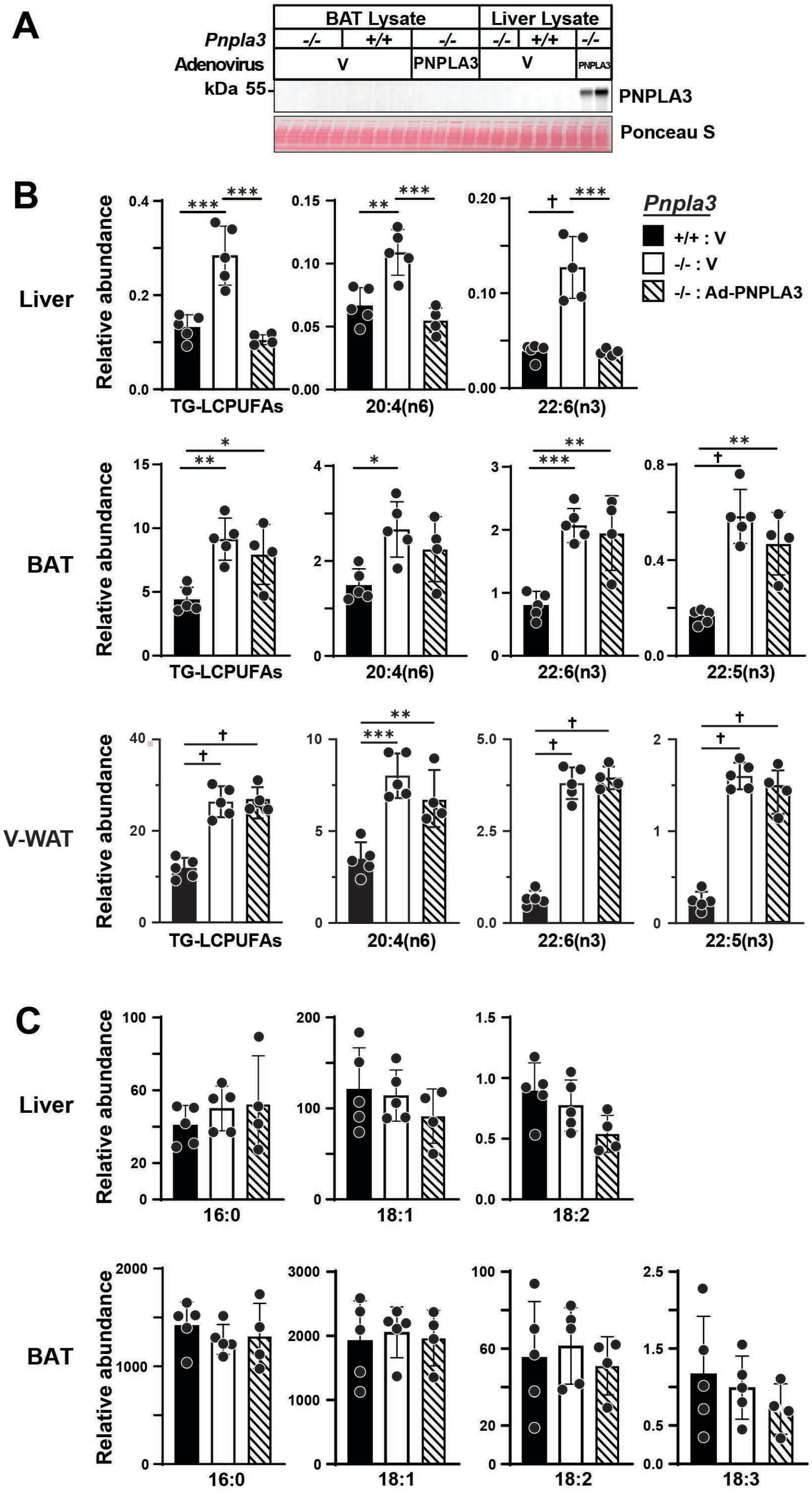


**Fig. S2. Composition of FAs in liver and adipose tissue of *Pnpla3*$^{-/-}$ and WT mice infected with vector alone (V) or Ad-PNPLA3-V5.** (A) Immunoblot of human PNPLA3 in lysates from

BAT and liver of mice (n = 4-5/group) infected with either vector alone (V) or Ad-PNPLA3-V5. (B) Relative abundance of TG-LCPUFAs from liver (upper), BAT (middle) and V-WAT (lower) from the same mice. (C) Relative abundance of other FAs (16:0, 18:1, 18:2, 18:3) in TG from liver (upper) and BAT (lower). Data represent mean ± SD. P values were determined by one-way ANOVA with Tukey's multiple comparisons test. * $P < 0.05$; ** $P < 0.01$; *** $P < 0.001$; †$P < 0.0001$.

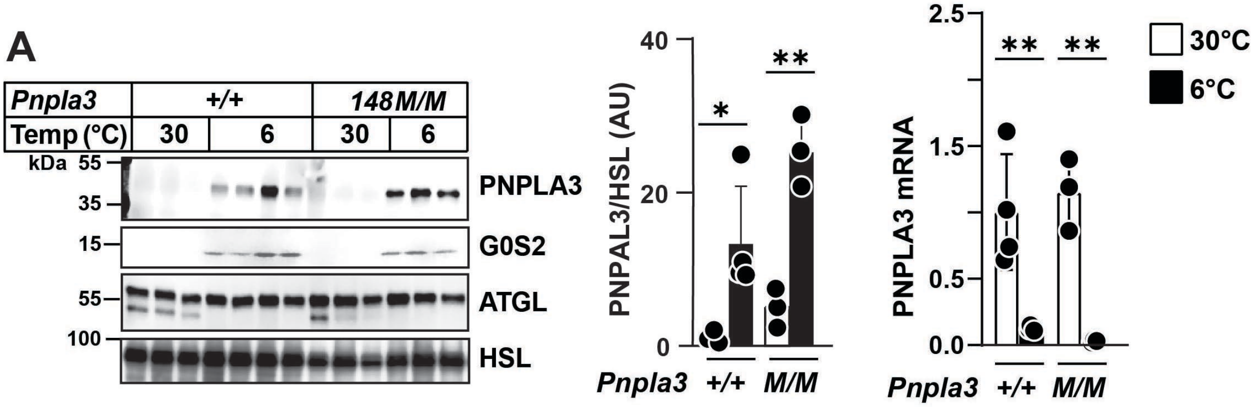


**Fig. S3. Cold exposure increases PNPLA3 protein while decreasing *Pnpla3* mRNA in BAT of WT and *Pnpla3$^{M/M}$* mice.** (A) Immunoblotting analysis of PNPLA3 protein and RT-qPCR analysis of *Pnpla3* mRNA in BAT from WT and *Pnpla3$^{M/M}$* mice. Mice (female, 6-10 weeks old; n = 3-4/group) were maintained at 30°C or 6°C for 12 h and fasted prior to sacrifice. PNPLA3 protein levels were quantified and normalized to HSL. Data are presented as mean ± SD. PNPLA3 protein and *Pnpla3* mRNA levels were compared between 30°C and 6°C within each genotype using Student's t-test; *P < 0.05; **P < 0.01.

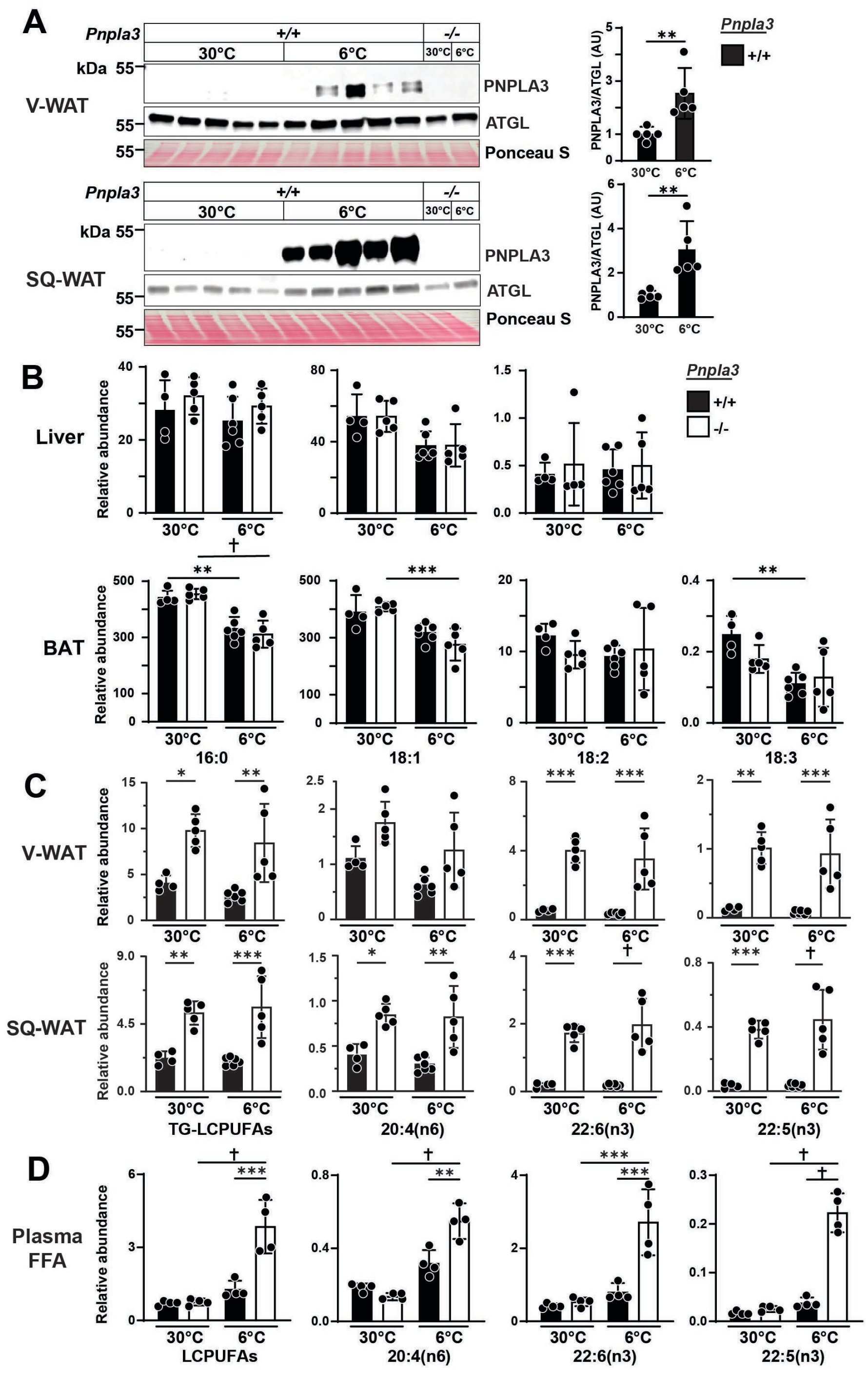

**Fig. S4. Composition of FAs in TGs of liver and adipose tissue of cold-exposed *Pnpla3*$^{-/-}$ and WT mice.** (A) Immunoblot analysis of PNPLA3 in fat cake fractions of V-WAT (upper) and SQ-WAT (lower) from mice (n = 5/group) maintained at 30°C or 6°C for one week on a HSD. (B) Relative abundance of other FAs (16:0, 18:1, 18:2, 18:3) in TG of liver (upper) and BAT (lower) from WT and *Pnpla3*$^{-/-}$ mice (n = 4-6/group) maintained at 30°C or 6°C for one week on a HSD. (C) Relative abundance of LCPUFAs in TG of V-WAT (upper) and SQ-WAT (lower) from the same mice. (D) Relative abundance of free LCPUFAs in plasma from WT and *Pnpla3*$^{-/-}$ mice (n = 4/group) maintained at 30°C or 6°C for one week on a HSD, and killed after 12 h fasting. Data represent mean ± SD. P values were determined by a Student's t-test (A) and by one-way ANOVA with Tukey's multiple comparisons test (B-D). * P < 0.05; ** P < 0.01; *** P < 0.001; †P < 0.0001.

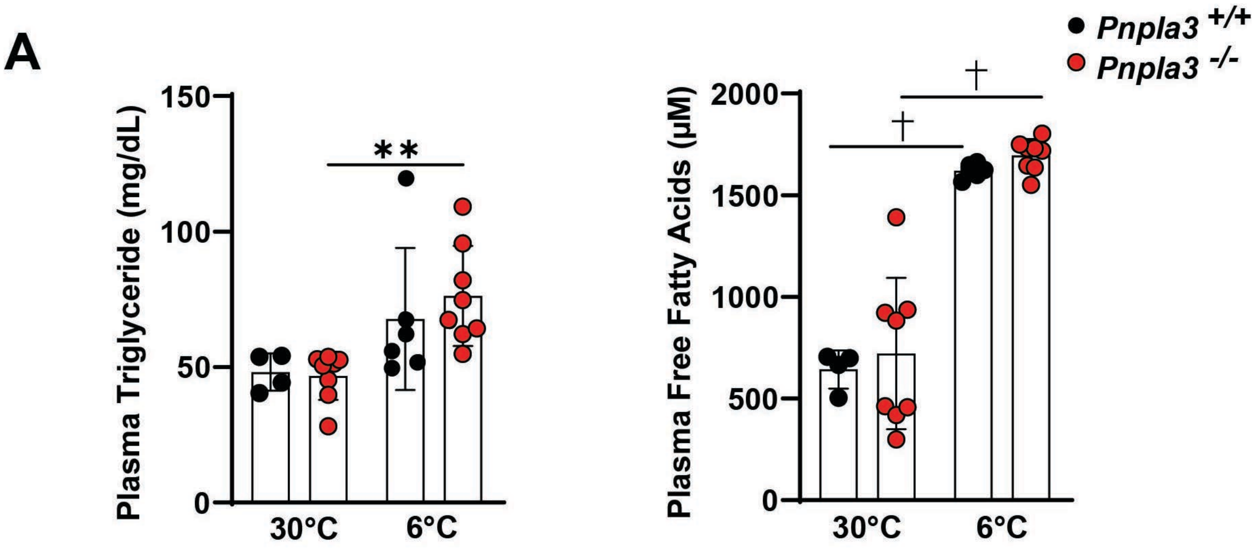


**Fig. S5. Effects of cold exposure on circulating TG and free fatty acid levels in WT and *Pnpla3*$^{-/-}$ mice.** (A) Plasma TG and free fatty acid levels in WT and *Pnpla3*$^{-/-}$ mice maintained at 30°C or 6°C. Mice (male, 11-15 weeks old; n = 4-8/group) were maintained on a high-sucrose diet and fasted for 12 h prior to sacrifice. Data are presented as mean ± SD. Statistical analysis was performed using one-way ANOVA with Tukey's multiple comparisons test. **P < 0.01; †P < 0.0001.

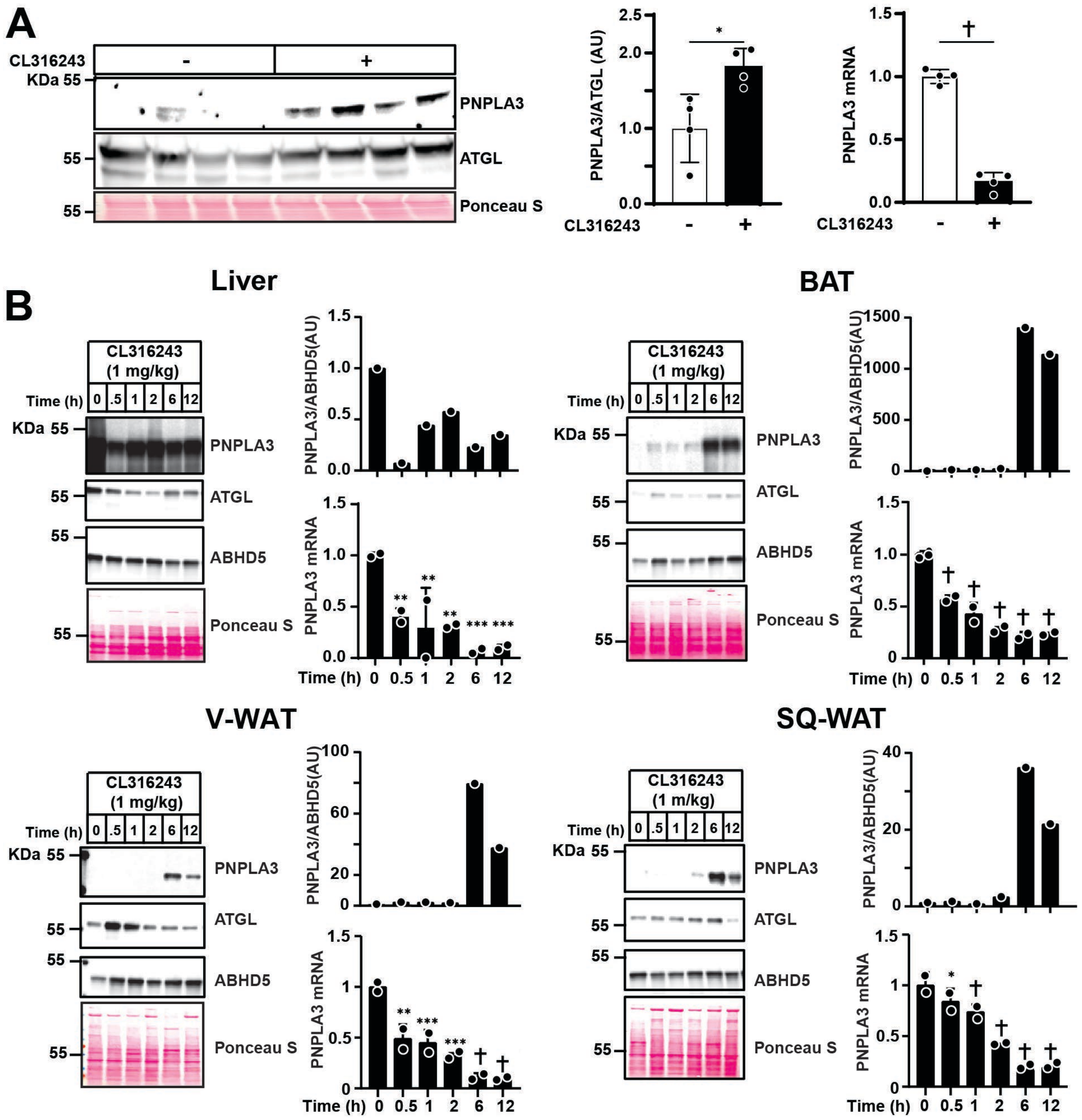


**Fig. S6. Effects of β3AR activation on PNPLA3 protein and mRNA expression in mouse liver and adipose depots.** (A) PNPLA3 protein abundance and *Pnpla3* mRNA levels in V-WAT from WT mice (n = 4/group) treated with β3AR agonist CL316243 or saline for 6 h. (B) PNPLA3 protein abundance and *Pnpla3* mRNA levels in liver, BAT, V-WAT, and SQ-WAT from WT mice (n = 2/group) treated with CL316243 for the indicated times. Data are presented as mean ± SD. P values were determined by Student's *t*-test (A) or by one-way ANOVA followed by Dunnett's multiple comparisons test (B); $^{*}P < 0.05$; $^{**}P < 0.01$; $^{***}P < 0.001$; $^{\dagger}P < 0.0001$.

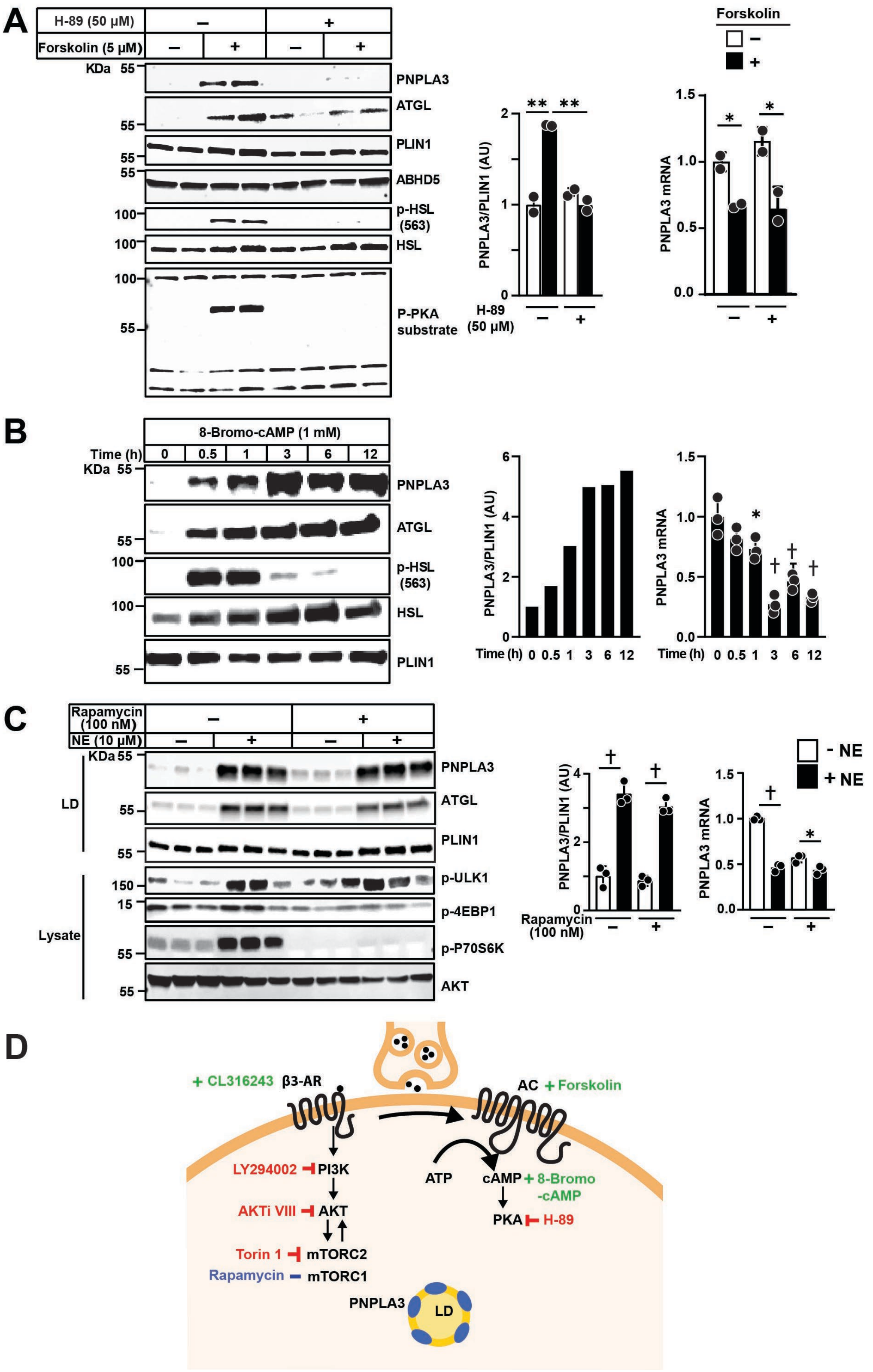


**Fig. S7. PNPLA3 expression in adipocytes is regulated by cAMP/PKA and PI3K/AKT signaling.** (A) Immunoblot analysis (left, middle) of PNPLA3 in LDs and RT-qPCR analysis (right)

of *Pnpla3* mRNA in 3T3-L1 cells treated with forskolin (5 μM) +/- H-89 (50 μM). (B) Immunoblot analysis of PNPLA3 in LD fractions (left, middle) and RT-qPCR of *Pnpla3* mRNA (right) in 3T3-L1 cells treated with 8-Bromo-cAMP (1 mM); cells were collected at the indicated time points. (C) Immunoblot analysis (left, middle) of PNPLA3 on LDs and RT-qPCR analysis (right) of *Pnpla3* mRNA in 3T3-L1 cells treated with norepinephrine (NE) (10 μM) +/- rapamycin (100 nM). (D) Schematic of signaling pathways: CL316243 activates β3-adrenergic receptors (β3AR), engaging the PI3K–AKT–mTORC axis (inhibited by LY294002, AKTi VIII and Torin 1) and, in parallel, the adenylyl cyclase–cAMP–PKA pathway (stimulated by forskolin or 8-Bromo-cAMP; blocked by H-89), with PNPLA3 on LDs as a downstream effector. Data represent mean ± SD (n = 2-3/group). P values were determined by one-way ANOVA followed by Tukey's multiple comparisons test (A, C) or by one-way ANOVA followed by Dunnett's multiple comparisons test (B). *P < 0.05; **P < 0.01; †P < 0.0001.

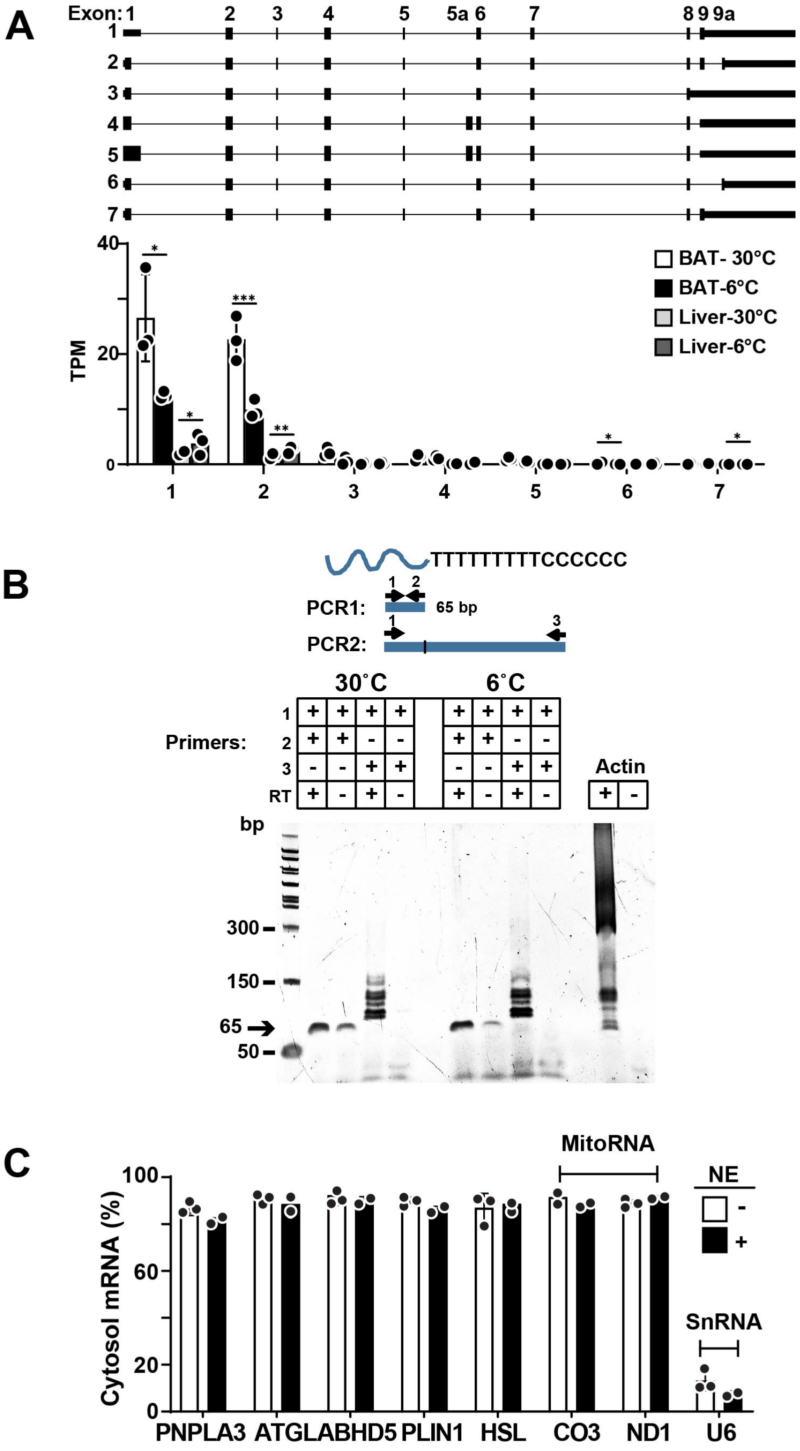


**Fig. S8. Post-transcriptional regulation of PNPLA3 expression in liver and BAT in response to temperature.** (A) RNA-seq analysis of *Pnpla3* transcripts in BAT and liver from WT mice (n = 3/group) housed at 30°C or 6°C for 12 h on a HSD. In both BAT and liver, seven *Pnpla3* isoforms were identified (upper); Transcript-level expression (TPM) values for each isoform are summarized in the bar chart (lower). (B) Polyacrylamide gel electrophoresis of poly(A)-tailed PCR products from BAT of WT mice housed at 30°C or 6°C for 12 h on HSD. (C) Partitioning of mRNAs between nuclear and cytosolic fractions of 3T3-L1 adipocytes +/- norepinephrine (NE) treatment for 6 h. Data represent mean ± SD (n = 2-3/group). P values were determined by an unpaired Student's *t*-test; $P < 0.05$.

### Supplementary Tables

### Table S1. Materials

| Reagent | Source | Identifier |
|---|---|---|
| Antibodies (Ab) | | |
| ABHD5 | Novus | H00051099-M01 |
| AKT | Cell Signaling Technology | 9272 |
| ATGL | Cell Signaling Technology | 2138 |
| Calnexin | Enzo Life Sciences | ADI-SPA-860-F |
| G0S2 | Proteintech | 12091-1-AP |
| HA | Biolegend | 901513 |
| HSL | Cell Signaling Technology | 18381 |
| P-4EBP1 | Cell Signaling Technology | 9451 |
| P-AKT473 | Cell Signaling Technology | 4060 |
| Phospho-(Ser/Thr) PKA | Cell Signaling Technology | 9621 |
| P-HSL563 | Cell Signaling Technology | 4139 |
| PLIN1 | Cell Signaling Technology | 3470 |
| PLIN2 | Abcam | ab108323 |
| P70S6K | Cell Signaling Technology | 9202 |
| P-P70S6K | Cell Signaling Technology | 9205 |
| PPARg | Cell Signaling Technology | 2435 |
| P-ULK1 | Cell Signaling Technology | 5869 |
| PNPLA3 (19A6) | Lab made | N/A |
| RPL22 | Santa Cruz | sc-522583 |

| | | |
|---|---|---|
| RPL7 | Novus Biologicals | NB100-2269 |
| RPS6 | Cell Signaling Technology | 2217 |
| Ubiquitin | Cell Signaling Technology | 43124 |
| V5 | Thermo Fisher Scientific | R960-25 |
| Peroxidase AffiniPure Goat Anti-Rabbit IgG (H+L) | Jackson Immunoresearch Laboratories | 111-035-144 |
| Peroxidase AffiniPure Donkey Anti-Mouse IgG (H+L) | Jackson Immunoresearch Laboratories | 715-035-150 |
| Rabbit TrueBlot®: Anti-Rabbit IgG HRP | Rockland Immunochemicals, Inc. | 18-8816-33 |
| Mouse TrueBlot® ULTRA: Anti-Mouse Ig HRP | Rockland Immunochemicals, Inc. | 18-8817-33 |
| Recombinant Viruses | | |
| Ad-RR5 | pShuttle; Clontech | |
| Ad-PNPLA3 (WT) | Lab made | |
| Buffers | | |
| 4X Laemmli Sample Buffer | Bio-Rad | 1610747 |
| 6X Laemmli Sample Buffer | Thermo Fisher Scientific | J61337.AC |
| TRIS Buffered Saline (TBS) | Sigma-Aldrich | T6664 |
| HEPES | Thermo Fisher Scientific | 15630080 |
| PBS, pH 7.4 | Gibco | 10010023 |
| RNA Gel Buffer (10X MOPS Buffer) | Fisher Scientific | 50-983-261 |

| | | |
|---|---|---|
| RIPA Lysis and Extraction Buffer | Thermo Fisher Scientific | 89900 |
| Cell Lines | | |
| 3T3-L1 murine fibroblasts | ATCC | CL-173 |
| Chemicals | | |
| Fetal Bovine Serum (FBS) | Millipore Sigma | F0926 |
| Bovine Serum Albumin (BSA), Cohn Fraction V | Avantor | J64944-22 |
| cOmplete Mini EDTA-free Protease Inhibitor Cocktail | Sigma-Aldrich | 11836170001 |
| Dimethyl Sulfoxide (DMSO) | Sigma-Aldrich | D2650 |
| Glycerol | Sigma-Aldrich | G9012 |
| Penicillin-Streptomycin | Corning | 30-002-Cl |
| MG132 | Peptide Institute, INC | 3178-v |
| LY294002 | Selleck Chemicals | S1105 |
| AKT inhibitor VIII | MedChemExpress | 612847-09-3 |
| Torin 1 | Selleck Chemicals | S2827 |
| Rapamycin | Sigma-Aldrich | 553211 |
| Forskolin | Millipore Sigma | F6886 |
| 8-Bromo-cAMP | Selleckchem | S7857 |
| H-89 dihydrochloride hydrate | Millipore Sigma | B1427 |
| Cycloheximide | Millipore Sigma | C7698 |
| Benzonase Nuclease | Millipore Sigma | E1014-25KU |

| | | |
|---|---|---|
| Insulin(cattle) | MedChemExpress | HY-P1156 |
| Dexamethasone | Millipore Sigma | D4902 |
| 3-Isobutyl-1-methylxanthine | Millipore Sigma | I7018 |
| Rosiglitazone | Millipore Sigma | R2408 |
| Tricine | Sigma | T0377 |
| Noradrenaline tartrate | Millipore Sigma | N1100000 |
| CL 316,243 hydrate | Millipore Sigma | C5976 |
| Sucrose, Ultrapure Bioreagent, J.T. Baker™ | Fisher Scientific | 02-004-331 |
| Sodium chloride | Sigma | S9625 |
| Potassium chloride | Millipore Sigma | P5405 |
| Magnesium chloride hexahydrate | Millipore Sigma | M9272 |
| Acetone, HPLC Grade, ≥ 99.5%, LabChem™ | Fisher Scientific | LC104254 |
| Ethyl Ether | Sigma-Aldrich | EX0185-4 |
| Sodium dodecyl sulfate solution | Millipore Sigma | 71736 |
| Urea | Sigma-Aldrich | U5378 |
| Enzymes | | |
| Pierce™ Trypsin Protease, MS Grade | Thermo Scientific | 90057 |
| DNase I (Lyophilized) | Promega | Z3585 |

| | | |
|---|---|---|
| SuperScript™ IV Reverse Transcriptase | Thermo Fisher Scientific | 18090010 |
| Kits | | |
| Free Fatty Acid Assay Kit | LSBio | LS-K170-100 |
| Glycerol Assay Kit | Millipore Sigma | MAK117 |
| Pierce BCA Protein Assay Kit | Thermo Fisher Scientific | 23224 |
| PARIS™ Kit | Invitrogen™ | AM1921 |
| Power SYBR Green PCR master Mix | Applied Biosystems | 4368708 |
| TaqMan reverse transcription reagents | Applied Biosystems | N8080234 |
| RNeasy Plus Universal Mini Kit | Qiagen | 73404 |
| SuperSignal™ West Pico Chemiluminescent Substrate | Thermo Fisher Scientific | 34580 |
| SuperSignal™ West Femto Maximum Sensitivity Substrate | Thermo Fisher Scientific | 34096 |
| TruSeq Stranded Total RNA | Illumina | 20020596 |
| Poly(A) Tail-Length Assay Kit | Invitrogen | 764551KT |
| Medium | | |
| DMEM High Glucose Medium | Corning | 10-013-CV |
| Oligonucleotides | | |
| qPCR primers: | Sequence | Supplier |

| | | |
|---|---|---|
| *Hprt* | CCTCATGGACTGATTATGGACAG; AATCCAGCAGGTCAGCAAAG | Integrated DNA Technologies IDT |
| *Cyclophilin B* | TGGAGAGCACCAAGACAGACA; TGCCGGAGTCGACAATGAT | Integrated DNA Technologies IDT |
| *Pnpla3* | CGAGGCGAGCGGTACGT; TGACACCGTGATGGTGGTTT | Integrated DNA Technologies IDT |
| *Abhd5* | AATGTGTCCCCTGCACTTACAA; GAACATCAGCGTCCATATTCTGTT | Integrated DNA Technologies IDT |
| *Atgl* | GAGAGAACGTCATCATATCCCACTT; CCACAGTACACCGGGATAAATGT | Integrated DNA Technologies IDT |
| *Ucp1* | ACTGCCACACCTCCAGTCATT; CTTTGCCTCACTCAGGATTGG | Integrated DNA Technologies IDT |
| *Plin1* | GGTGAGCGGGACCTGTGA; TTCTCATAGGCATTGCACACAGA | Integrated DNA Technologies IDT |
| *U6* | GTGCTCGCTTCGGCAGC; | Integrated DNA |

| | | |
|---|---|---|
| | AAAAATATGGAACGCTTCACGAAT | Technologies IDT |
| *Hsl* | GGAGCACTACAAACGCAACGA; TCGGCCACCGGTAAAGAG | Integrated DNA Technologies IDT |
| *Co3* | CCAAGGCCACCACACTCCTA; GGTCAGCAGCCTCCTAGATCA | Integrated DNA Technologies IDT |
| *Nd1* | GCTTTACGAGCCGTAGCCCA; GGGTCAGGCTGGCAGAAGTAA | Integrated DNA Technologies IDT |
| Software and Algorithms | | |
| Image Studio Lite | LI-COR | v5.2 (version) |
| Prism 10 | GraphPad | 10.2.3 (version) |
| Other | | |
| Nitrocellulose Membrane | Bio-Rad | 1620168 |
| 4–15% Criterion™ TGX™ Precast Midi Protein Gel, 26 well, 15 µl | Bio-Rad | 5671085 |
| 4–15% Criterion™ TGX™ Precast Midi Protein Gel, 18 well, 30 µl | Bio-Rad | 5671084 |
| 4–15% Mini-PROTEAN® | Bio-Rad | 4561084 |

| | | |
|---|---|---|
| TGX™ Precast Protein Gels, 10-well, 50 µl | | |
| 5% Criterion™ TBE Polyacrylamide Gel, 12+2 well, 45 µl 3450047 | Bio-Rad | 3450047 |
| InstantBlue® Coomassie Protein Stain (ISB1L) | Abcam | ab119211 |
| Pierce™ Protein G Magnetic Beads | Thermo Fisher Scientific | 88848 |
| RNasin® Plus Ribonuclease Inhibitor | Promega | N2615 |
| Heavy-isotope labeled peptide-DGLQESLPDNVHQVISGK (aa 96–113) | 21st Century Biochemicals | N/A |
| Heavy-isotope labeled peptide-YVDGGVSDNVPVLDAK (aa 163–179) | 21st Century Biochemicals | N/A |
| Heavy-isotope labeled peptide-STNFFHVNITNLSLR (aa 188–213) | 21st Century Biochemicals | N/A |
